# Cryo-EM structure of DNA-unbound human MCM2–7 complex reveals new disease-relevant regulation

**DOI:** 10.1101/2025.05.31.656953

**Authors:** Yusong Liu, Mengquan Yang, Ping Lu, Haishan Gao, Maozhou He, Yitao Wang, Ao Qi, Ting Cao, Qiuqin Zhang, Shutao Qi, Yigong Shi, Hongtao Yu

## Abstract

Chromatin loading of the hexameric replicative helicase MCM2–7 complex requires coordinated interactions with the origin recognition complex (ORC), CDC6, and CDT1. MCM2–7 not bound to DNA forms a single hexamer (SH) with an open DNA entry gate. Two MCM2–7 SHs are loaded sequentially to form the double hexamer (DH) that encircles the DNA duplex. Activated MCM2–7 then unwinds DNA and initiates DNA replication. Our cryo-electron microscopy analyses show that a fraction of human MCM2–7 without DNA exists as DH. Unexpectedly, we find that the MCM3 winged helix domain (WHD) docks on MCM2 in both DNA-free DH and SH, creating a safety latch across the DNA entry gate to block DNA entry into the central channel. The safety latch can be opened by ORC-CDC6 binding. Disrupting this latch by designed or human disease-related mutations of MCM3 causes replication defects and DNA damage checkpoint activation. Our findings uncover a new regulated step in MCM2–7 loading with implications for human diseases.

## Introduction

Faithful replication of genomic DNA ensures genome stability. DNA replication is restricted to once-and-only-once in each cell division cycle. The origin recognition complex (ORC), which consists of six subunits (ORC1–6), binds to replication origins throughout the cell cycle.^1–3^ In G1, ORC, CDC6, and CDT1 act together to load the hexameric replicative helicase MCM2–7 onto chromatin.^1–3^ In S phase, MCM2–7 is activated by phosphorylation and the binding of CDC45 and GINS, forming the active CDC45-MCM2–7-GINS (CMG) helicase, which unwinds DNA, nucleates replisome formation, and initiates DNA replication.^4^ MCM2–7 loading is permitted in G1 when the overall cellular CDK activities are low whereas MCM2–7 activation is only possible in S phase when CDK activities are high.^1^ MCM2–7 reloading cannot occur in G2/M when the CDK activities are high. The differential regulation of MCM2–7 loading and activation by CDK activities ensures the proper coordination of DNA replication with cell division.

*In vitro* reconstitution, cryo-electron microscopy (cryo-EM), and single-molecule biophysical experiments of the assembly of the pre-replication complex (pre-RC; particularly that of the budding yeast) have provided key insight into the molecular mechanisms of MCM2-7 loading and activation.^5^ In the absence of DNA, free MCM2–7 exists as a single hexamer (SH) with an open DNA entry gate located between MCM2 and MCM5.^6–9^ The DNA-binding central channel of MCM2–7 is, however, occupied by winged helix domains (WHDs) of MCM4 and MCM5.^10,11^ During pre-RC assembly, CDC6 joins the DNA-bound ORC to form a functional ATPase.^12–14^ ORC-CDC6 then recruits and loads the CDT1-stabilized MCM2–7 SH to form the ORC-CDC6-CDT1-MCM2–7 (OCCM) complex on DNA.^8,10,15,16^ After CDC6 is released, ORC switches positions relative to the loaded MCM2–7 SH to form the MCM2–7-ORC (MO) complex and loads the second CDT1-stabilized MCM2–7 SH onto DNA.^17–19^ The two loaded MCM2–7 SHs form a head-to-head double hexamer (DH) to encircle the DNA duplex after the release of ORC-CDC6 and CDT1.^17^

While the overall framework of MCM2–7 loading and regulation is conserved among eukaryotes, there are species-specific features in this process. For example, yeast MCM2–7 is stably bound by CDT1 whereas human MCM2–7 by itself does not bind CDT1 tightly.^9,20,21^ Recent evidence also suggests that human MCM2–7 can be loaded onto DNA by both ORC6-dependent and -independent pathways *in vitro.*^22,23^ Yeast MCM2–7 DH encircles an undistorted DNA duplex in its central channel, but human MCM2–7 DH melts the DNA duplex at its hexamer junction.^6,22–24^

Mutations of human MCM2–7 have been linked to human developmental diseases, such as the Meier-Gorlin syndrome (MGORS), which is characterized by dwarfism and other abnormalities.^25,26^ To better understand the species-specific features of MCM2–7 loading and the detrimental effects of disease-causing mutations, we determined the cryo-EM structures of the endogenous MCM2–7 complex purified from human cells and the recombinant human MCM2–7 expressed in insect cells. We found that a fraction of human MCM2–7 complex formed DHs in the absence of DNA. Cryo-EM structures of DNA-free MCM2–7 SH and DHs showed that MCM3 WHD docked onto the C-terminal ATPase lobe of MCM2, forming a safety latch to prevent DNA from accessing the central channel of MCM2–7. Interestingly, an MCM3 mutation linked to MGORS strengthened this latch and inhibited MCM2–7 loading in human cells. The same mutation also caused DNA replication defects and activated G2/M DNA damage checkpoint. Our study thus revealed a novel regulatory mechanism in MCM2–7 loading with disease implications.

## Results

### Cryo-EM structures of DNA-unbound human MCM2–7 SH and DH

To purify the endogenous human MCM2–7 complex, we introduced three tandem copies of the FLAG tag and two copies of the StrepII tag into the *MCM4* loci, or three tandem copies of the Flag tag into the *MCM7* loci, in human HEK293 cells using the CRISPR-Cas9 technology (Figures 1A, S1A, and S1B). The endogenous DNA-unbound MCM2–7 complex was purified from MCM4-FLAG-StrepII or MCM7-FLAG HEK293 cells using anti-FLAG antibody beads followed by density gradient centrifugation (Figures S1C and S1D). Consistent with a previous study,^9^ both negative-stain EM and cryo-EM showed that, while the majority of MCM2–7 particles belonged to single hexamers (SH), a small fraction of MCM2–7 formed double hexamers (DH) (Figures S1E-G). The overall conformation of SH was very similar to that of the DNA-unbound yeast and human MCM2–7 SH.^7,8,23^ The structure of each SH in the DH was also virtually identical to that of the SH alone. We focus our description on the structure of DNA-unbound human MCM2–7 DH, as it has a higher resolution and has not been extensively reported.

**Figure 1.**
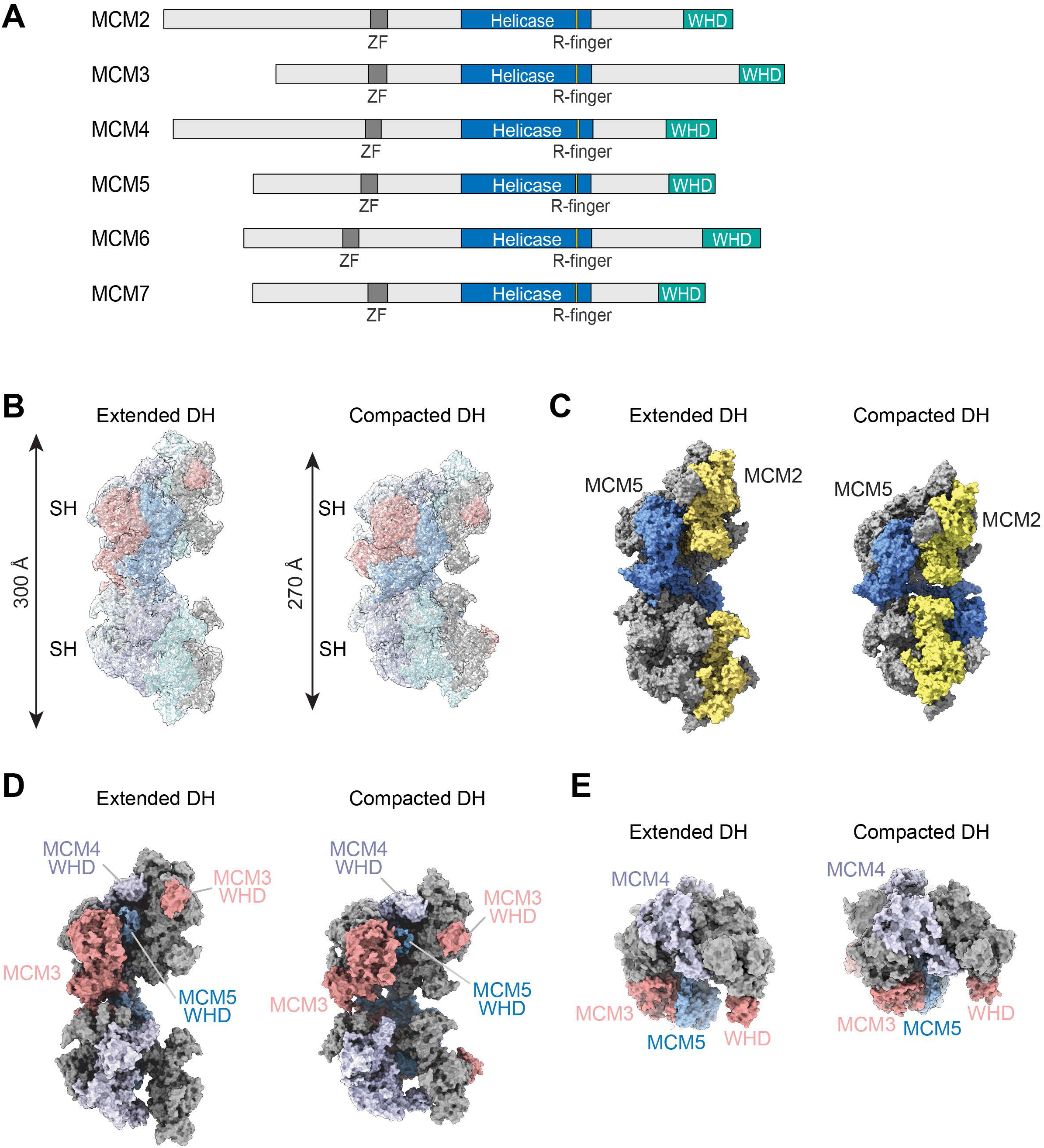
Overall architecture of DNA-unbound human MCM2–7 double-hexamer. (A) Schematic illustration of the domain organization of human MCM2–7 subunits. ZF, zinc finger; R-finger, arginine finger; WHD, winged helix domain. (B) Side views of the segmented cryo-EM density maps of the extended and compacted conformers of DNA-unbound human MCM2–7 double hexamer (DH), superimposed with the atomic models. (C) Side views of the two MCM2–7 DH conformers showing the open DNA-entry gate between MCM2 (yellow) and MCM5 (blue). (D) Side views of two MCM2–7 DH conformers with two subunits removed to display the DNA-binding central channel and the WHDs of MCM4 and MCM5, which occupy the central channel. (E) Top views of the two MCM2–7 DH conformers showing the DNA-entry gate.

We determined the cryo-EM structures of two DH conformers––the extended and compacted DHs––with resolutions of 3.8 Å and 3.9 Å, respectively (Figures 1B, S2, S3, and S4 and Table S1). Both conformers are head-to-head stacked DHs that consist of two symmetric SHs with their N-terminal tiers forming the dimerization interface (Figure 1B). The lengths of the extended and compacted DHs are about 300 Å and 270 Å, respectively. The inter-SH dimerization interfaces in the two conformers are similar. In both conformers, the N-terminal extension of MCM5 (residues 1-21) of one SH interacts extensively with the helicase domain of MCM7 in the other (Figures S5A and S5B). The zinc fingers (ZFs) of MCM3 and MCM5 from one SH pack against those from the other (Figure S5C). There are also important differences between the two interfaces. For example, the ZFs of the two MCM5 molecules directly contact each other in the extended conformer, but not in the compacted conformer (Figure S5C). The underlying reason of the conformational differences between the two conformers is unknown, but may be caused by differences in bound nucleotides (ATP or ADP) of MCM subunits.

In both DH conformers, the two symmetric SHs adopt a left-handed spiral conformation with an open DNA-entry gate between MCM2 and MCM5 (Figure 1C). The DNA entry gates of the two SHs in the extended and compacted DHs are not aligned and are related by rotation angles of about 90° and 70°, respectively. The DNA-binding channels and the central axes of the SHs are also not aligned in both DH conformers (Figures 1C and S5D). The offset between the two SH axes in these DNA-free DHs is larger than that in the DNA-bound human MCM2–7 DH.^24^ The central channel of DHs has a diameter larger than 20 Å, which is wide enough to accommodate double-stranded DNA (dsDNA) (Figure 1D). The C-terminal WHDs from MCM4 and MCM5, however, occupy this central channel, preventing DNA from binding (Figures 1D and 1E). Overall, the spiral, open-gated architecture of our DNA-free SH resembles the cryo-EM structure of yeast MCM2–7 in complex with CDT1 before loading and represents the pre-loading state of MCM2–7 SH.^5^ Comparison with the DNA-bound yeast and human MCM2–7 indicates that the release of MCM4 and MCM5 WHDs from the central channel and the DNA-entry gate closure are needed to load MCM2–7 SH onto DNA.

MCM2–7 DH formation is a critical step in DNA replication initiation in eukaryotes and is generally thought to only occur after the sequential loading of two SHs onto DNA. Our finding that human MCM2–7 without DNA can form an open-gated DH in the preloading state is thus surprising. Future experiments are needed to explore the functional significance of the DNA-free MCM2–7 DH.

### MCM3 WHD forms a latch that guards the DNA entry gate

Focused refinement revealed an extra cryo-EM density attached to the C-terminal ATPase lobe of MCM2 in both SHs and DHs (Figure S6A). After extensive density analysis and model building, we assigned this density to the C-terminal winged helix domain (WHD) of human MCM3. Map-to-model auto-building by ModelAngelo could also fit MCM3 WHD into the extra density (Figure S6B).^27^ WHD is commonly found in chromatin-bound proteins and participates in protein-DNA and protein-protein interactions.^28^ All MCM2–7 subunits contain a WHD (Figure 1A). These WHDs have important roles in binding ORC and DNA. MCM3 WHD has previously been shown to interact with DNA-bound ORC-CDC6 in yeast.^29^ In our MCM2–7 complex structures, MCM3 WHD packs against the C-terminal ATPase lobe of MCM2, forming a conserved, hydrophobic interface (Figures 2A and 2B). The MCM3 WHD binding surface on MCM2 overlaps with its MCM5-binding interface in the DNA-bound SH or DH (Figures 2C and 2D).^23,24^ Thus, MCM3 WHD needs to be released from MCM2 to allow the closure of the MCM2–7 hexameric helicase ring and DNA entrapment.

**Figure 2.**
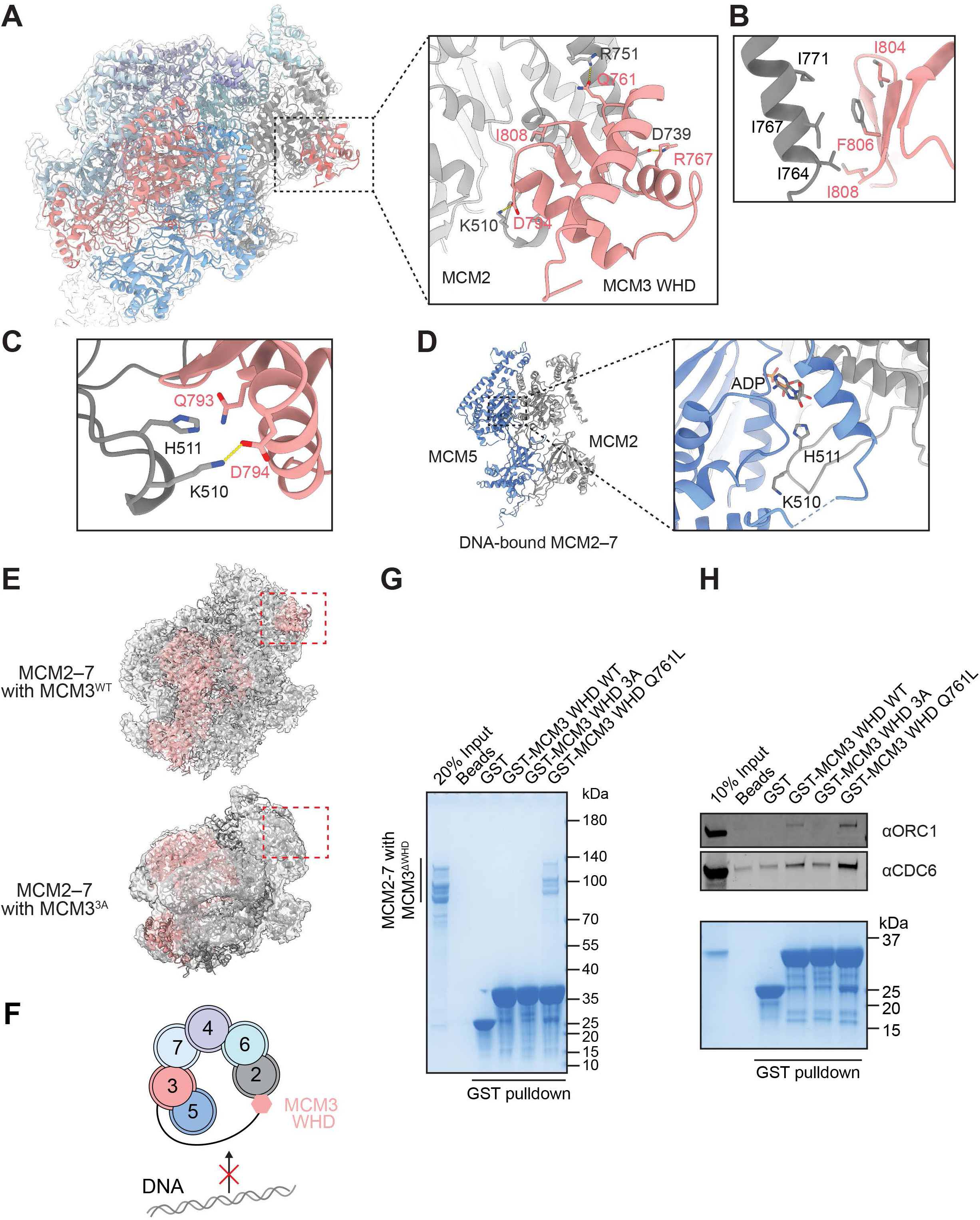
MCM3 WHD binds to MCM2 in DNA-unbound MCM2–7 DH. (A) Cryo-EM density map of an MCM2–7 single hexamer (SH) superimposed with the atomic model, with a close-up view of the MCM3 WHD-MCM2 interface. (B and C) Close-up views of key interacting residues at the MCM3 WHD–MCM2 interface. (D) The closed DNA entry gate in DNA-bound MCM2–7 DH (PDB ID: 7W1Y) with only MCM2 and MCM5 shown. A close-up view of the MCM2-MCM5 interface is shown on the right. (E) Cryo-EM density maps of recombinant human MCM2–7 SH that contains either MCM3 wild type (WT; top) or the MCM3 3A mutant (bottom). The density of MCM3 WHD (top) and its corresponding location (bottom) are boxed by red dashed lines. (F) Schematic diagram depicting the closed MCM3 WHD latch, which is expected to block DNA entry. (G) Binding between GST-MCM3 WHD proteins (WT, 3A, and Q761L) and the recombinant MCM2–7 complex containing MCM3 with its WHD deleted (MCM3^ΔWHD^). The input proteins and proteins bound to GST beads were analyzed by SDS-PAGE and Coomassie staining. The experiment was repeated three times with similar results. (H) Binding between GST-MCM3 WHD proteins (WT, 3A, and Q761L) and the ORC1-6–CDC6 complex. The input proteins and proteins bound to GST beads were analyzed by SDS-PAGE followed by Coomassie staining (bottom panel) and immunoblotting with anti-ORC1 and anti-CDC6 antibodies (top panels). The experiment was repeated three times with similar results.

To validate the MCM3 WHD–MCM2 interface and test whether MCM3 WHD was required to keep the DNA entry gate open, we expressed human MCM2–7 WT and MCM3^3A^ mutant (with three interface residues––I804, F806, and I808––of MCM3 WHD mutated to alanine) in insect cells, purified the resulting complexes, and determined their SH cryo-EM structures (Figure 2E). The cryo-EM map of MCM2–7 SH with the MCM3^3A^ mutation was virtually identical to that of MCM2–7 WT, except that the extra density belonging to MCM3 WHD was missing. Notably, even without MCM3 WHD bound to MCM2, MCM5 did not prematurely interact with MCM2 and the DNA entry gate between MCM2 and MCM5 remained open. Thus, the MCM3^3A^ mutation disrupted the interaction between MCM3 WHD and MCM2, validating the observed interface. It also indicates that the MCM3 WHD–MCM2 interaction was not required to maintain the open DNA-entry gate of MCM2–7 SH.

The MCM3 helicase domain is located on one side of the DNA entry gate of MCM2–7 while MCM3 WHD via its interaction with MCM2 is located on the other side (Figure 2A). The covalent linker connecting MCM3 WHD and its helicase domain is expected to straddle the DNA entry gate, even though we did not observe density belonging to the linker. This linker and MCM3 WHD thus form a safety latch that guards the DNA entry gate (Figure 2F), reminiscent of the chain locks found on some hotel room and apartment doors. This latch needs to be opened (i.e. MCM3 WHD needs to detach from MCM2) for DNA to enter the central channel of MCM2– 7.

The MCM3 Q761L mutation has been linked to the human developmental disorder MGORS.^25^ The functional consequences of this mutation on DNA replication have not been explored, however. Interestingly, MCM3 Q761 is located at the MCM3 WHD–MCM2 interface, contacting hydrophobic residues V743, A744, and Y747 of MCM2 (Figure S6C). To test whether MCM3 Q761L mutation affected the MCM3 WHD–MCM2 interaction, we performed GST pulldown assay between GST-MCM3 WHD and the recombinant MCM2–7 complex with MCM3 WHD deleted (MCM3^ΔWHD^) (Figure 2G). We could not detect an interaction between MCM3 WHD wild type (WHD^WT^) and MCM2–7^ΔWHD^, indicating that the affinity between MCM3 WHD and MCM2 was weak and that this interaction could only occur within the same complex. Interestingly, MCM3 WHD^Q761L^ efficiently pulled down MCM2–7 with MCM3^ΔWHD^, indicating that the Q761L mutation strengthened the interaction between MCM3 WHD and MCM2. This increased affinity of the Q761L mutant is consistent with the hydrophobic nature of Q761-binding pocket on MCM2, as leucine is more hydrophobic than glutamine. Thus, MCM3 WHD^Q761L^ forms a stronger latch as compared to the wild type.

In yeast, MCM3 WHD interacts with ORC-CDC6 and mediates pre-RC formation.^10,29^ Consistent with this finding, GST-tagged human MCM3 WHD pulled down both human ORC and CDC6 (Figure 2H), indicating that human MCM3 WHD interacted with ORC-CDC6. By contrast, MCM3 WHD^3A^ showed no detectable binding to ORC and only background binding to CDC6. Thus, the MCM3^WHD^ surface involved in MCM2 binding is also required for ORC-CDC6 binding. Recruitment of MCM2–7 to chromatin by ORC-CDC6 is expected to open the MCM3^WHD^ latch. Interestingly, MCM3 WHD^Q761L^ showed stronger ORC-CDC6 binding than WHD^WT^ (Figure 2H). Recent human OCCM structure demonstrated that Q761 bound to a hydrophobic pocket at the ORC2–CDC6 interface (Figure S6D),^22^ explaining why the glutamine-to-leucine substitution increased the affinity between MCM3 WHD and ORC-CDC6. The Q761L mutation thus strengthens both the MCM3 WHD–MCM2 interaction and the binding of MCM3 WHD to ORC-CDC6. It might alter the relative strengths of these two competing binding events and affect the latch opening by ORC-CDC6 and hence the recruitment of MCM2–7 to chromatin.

### MCM3 WHD mutations induce ATR-CHK1-dependent G2 arrest

To probe the cellular function of the MCM3 WHD latch, we tested the effects of MCM3 WHD^3A^ and WHD^Q761L^ on cell cycle progression in human cells. By using CRISPR-Cas9 knock-in (KI) technology, we tagged both alleles of the endogenous *MCM3* gene with the degradation tag (dTAG; FKBP12^F36V^) in HEK293FT cells (Figures 3A and S7A). The dTAG-MCM3 fusion protein could be induced for degradation upon the addition of the chemical ligand dTAG-13.^30^ We further transfected dTAG-MCM3 KI cells with plasmids encoding HA-MCM3 wild type (WT), WHD^3A^, or WHD^Q761L^. These HA-MCM3 transgenes produced HA-MCM3 proteins at levels comparable to that of the endogenous MCM3 (Figure 3B). Induced degradation of the endogenous MCM3 by dTAG-13 resulted in cells that expressed HA-MCM3 WT or the WHD mutants as the main MCM3 species.

**Figure 3.**
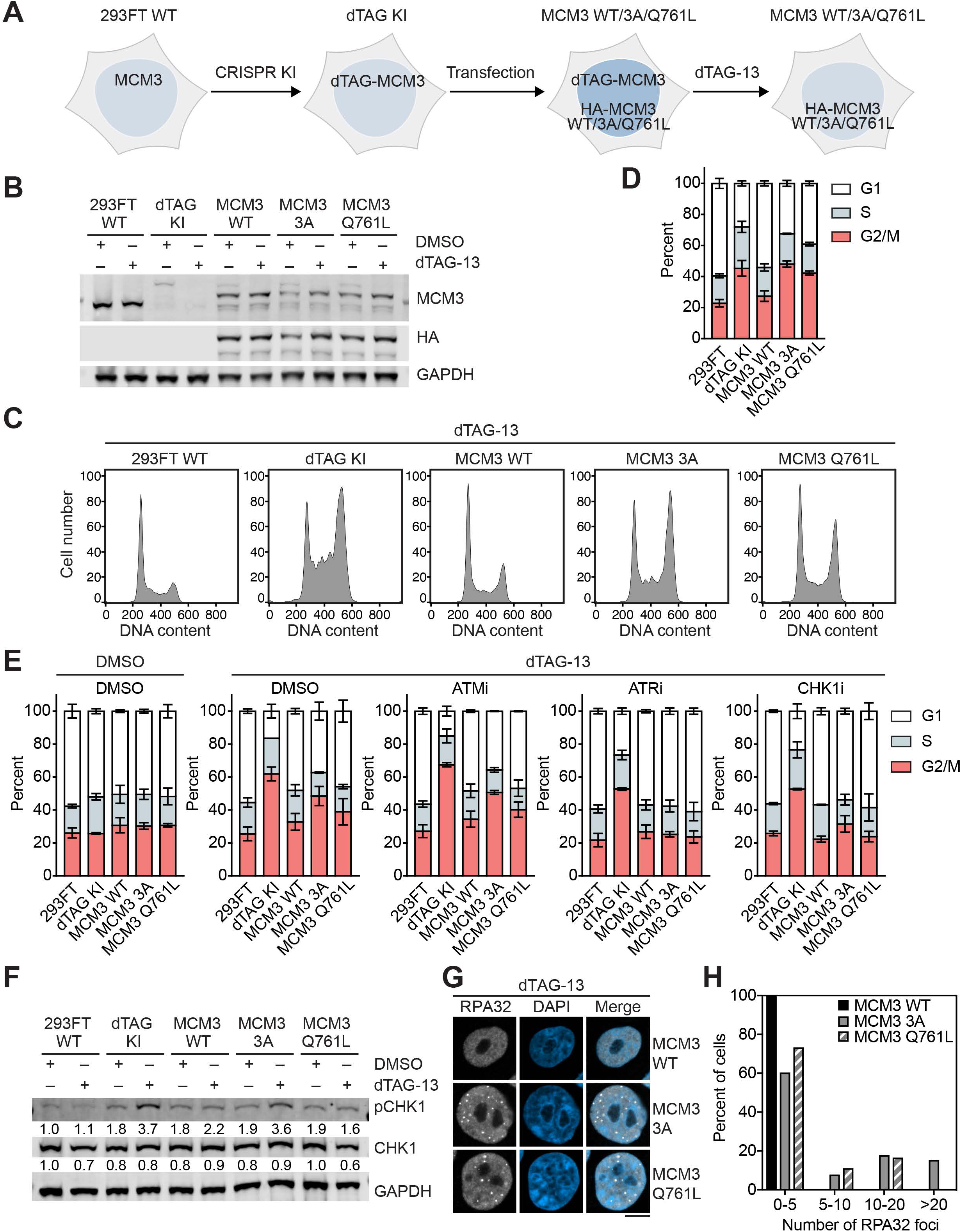
MCM3 WHD mutations induce ATR-CHK1-dependent G2 arrest in human cells. (A) Experimental strategy to deplete the endogenous MCM3 protein and ectopically express HA-MCM3 wild type (WT), 3A, or Q761L in 293FT cells. 293FT WT, the parental cell line without gene editing; dTAG KI, 293FT cells with dTAG knock-in (KI) at the endogenous *MCM3* loci; MCM3 WT, dTAG KI 293FT cells stably expressing HA-MCM3 wild type; MCM3 3A, dTAG KI 293FT cells stably expressing HA-MCM3 3A; MCM3 Q761L, dTAG KI 293FT cells stably expressing HA-MCM3 Q761L. (B) Immunoblots of whole-cell extracts of the indicated 293FT cells described in (A) with or without dTAG-13 (1 µM) treatment for 16 h. GAPDH was used as the loading control. 293FT WT, the parental cell line without gene editing; dTAG KI, 293FT cells with dTAG knock-in (KI) at the endogenous *MCM3* loci; MCM3 WT, dTAG KI 293FT cells stably expressing HA-MCM3 wild type; MCM3 3A, dTAG KI 293FT cells stably expressing HA-MCM3 3A; MCM3 Q761L, dTAG KI 293FT cells stably expressing HA-MCM3 Q761L. (C) Representative flow cytometry (FACS) profiles of the indicated 293FT cells treated with dTAG-13. (D) Quantification of cell cycle distributions of the indicated 293FT cells determined by FACS (C). Mean ± SD are shown; n = 4 biological replicates. (E) Cell cycle distributions of the indicated 293FT cells treated with DMSO or the indicated inhibitors with or without dTAG-13. Mean ± SD; n = 2 biological replicates. (F) Immunoblots of whole-cell extracts of the indicated 293FT cells with or without dTAG-13 (1 µM) treatment for 16 h. The relative intensities of the total CHK1 and pCHK1 (phospho-S345 CHK1) signals are shown below. (G) Representative images of the indicated 293FT cells treated with dTAG-13 for 16 h and stained with anti-RPA32 (white) and DAPI (blue). Scale bar, 10 µm. (H) Quantification of the percentage of cells in (G) that have the indicated numbers of RPA32 foci per cell.

As evidenced by flow cytometry (FACS) analysis of DNA content, depletion of the endogenous MCM3 for 16 hours caused the accumulation of cells in G2/M phases of the cell cycle (Figures 3C and 3D). FACS analysis with the MPM2 antibody that stained mitotic phospho-proteins further showed that these cells had a lower mitotic index as compared to cells without MCM3 depletion (Figure S7B), indicating that MCM3 depletion caused G2 arrest. Ectopic expression of HA-MCM3 WT rescued this G2 arrest phenotype and restored the normal cell cycle profile and mitotic index. By contrast, expression of HA-MCM3 3A or Q761L mutants did not fully rescue the G2 arrest phenotype, indicating that these mutants were functionally deficient and could not perform all cellular functions MCM3.

Because of the essential roles of MCM2–7 in DNA replication, complete inactivation of MCM3 is expected to block cells in S phase. However, it is well known that there is an excess amount of MCM2–7 in human cells, compared to the number of active replication forks.^31,32^ In each S phase, only a fraction of MCM2–7 DHs is activated to become CMG helicases.^33,34^ It is thus exceedingly difficult to deplete MCM2–7 to sufficiently low levels that actually block DNA replication. We hypothesized that partial depletion of MCM3 and expression of MCM3 3A and Q761L mutants could replicate most genomic DNA but caused replication stress, which in turn activated the G2/M DNA damage checkpoint, resulting in G2 arrest. Replication stress is more reliant on the ATR-CHK1 branch of the G2/M checkpoint whereas DNA double-stranded breaks preferably activate the ATM-CHK2 pathway.^35^ Indeed, chemical inhibitors of ATR and CHK1 reduced the G2 arrest of cells with MCM3 depletion and expression of HA-MCM3 3A and Q761L mutants (Figures 3E and S8). In contrast, an ATM inhibitor had no effect. Furthermore, after the depletion of the endogenous MCM3, cells expressing HA-MCM3 3A and Q761L had elevated levels of phospho-CHK1, consistent with its activation (Figure 3F). The replication protein A (RPA) complex binds to single-stranded DNA (ssDNA) at the replication fork, serving as the platform for ATR recruitment and activation during replication stress.^35^ Immunofluorescence staining against RPA32 showed increased RPA foci in HA-MCM3 3A- and Q761L-expressing cells with MCM3 depletion, indicative of replication stress in these cells (Figures 3G and 3H). Therefore, the MCM3 3A and Q761L mutations lead to replication stress-induced ATR-CHK1-dependent G2 arrest. Weakening or strengthening the MCM3 WHD latch thus causes replication defects, demonstrating its functional importance.

### MCM3 WHD mutations reduce chromatin-bound MCM levels

We next explored the mechanism by which MCM3 WHD contributed to DNA replication. Because MCM3 WHD interacts with ORC-CDC6, we examined whether MCM3 3A and Q761L mutations affected chromatin levels of MCM2–7. We performed chromatin fractionation assays to quantify chromatin-bound MCM2–7. While the total cellular levels of HA-MCM3 WT, 3A, and Q761L were similar, the levels of chromatin-bound HA-MCM3 3A and Q761L were lower than that of HA-MCM3 WT with or without the depletion of the endogenous MCM3 (Figure 4A). The levels of chromatin-bound MCM2, another subunit of MCM2–7, were also reduced in cells expressing HA-MCM3 3A or Q761L and with the depletion of endogenous MCM3. Immunofluorescence staining against HA confirmed the reduced chromatin association of HA-MCM3 3A and Q761L (Figures 4B-4E). The staining of chromatin-bound MCM5 was also reduced in 3A- or Q761L-expressing cells with MCM3 depletion (Figures 4F-4I). Thus, the 3A and Q761L mutations reduced the levels of the MCM2–7 complex on chromatin.

**Figure 4.**
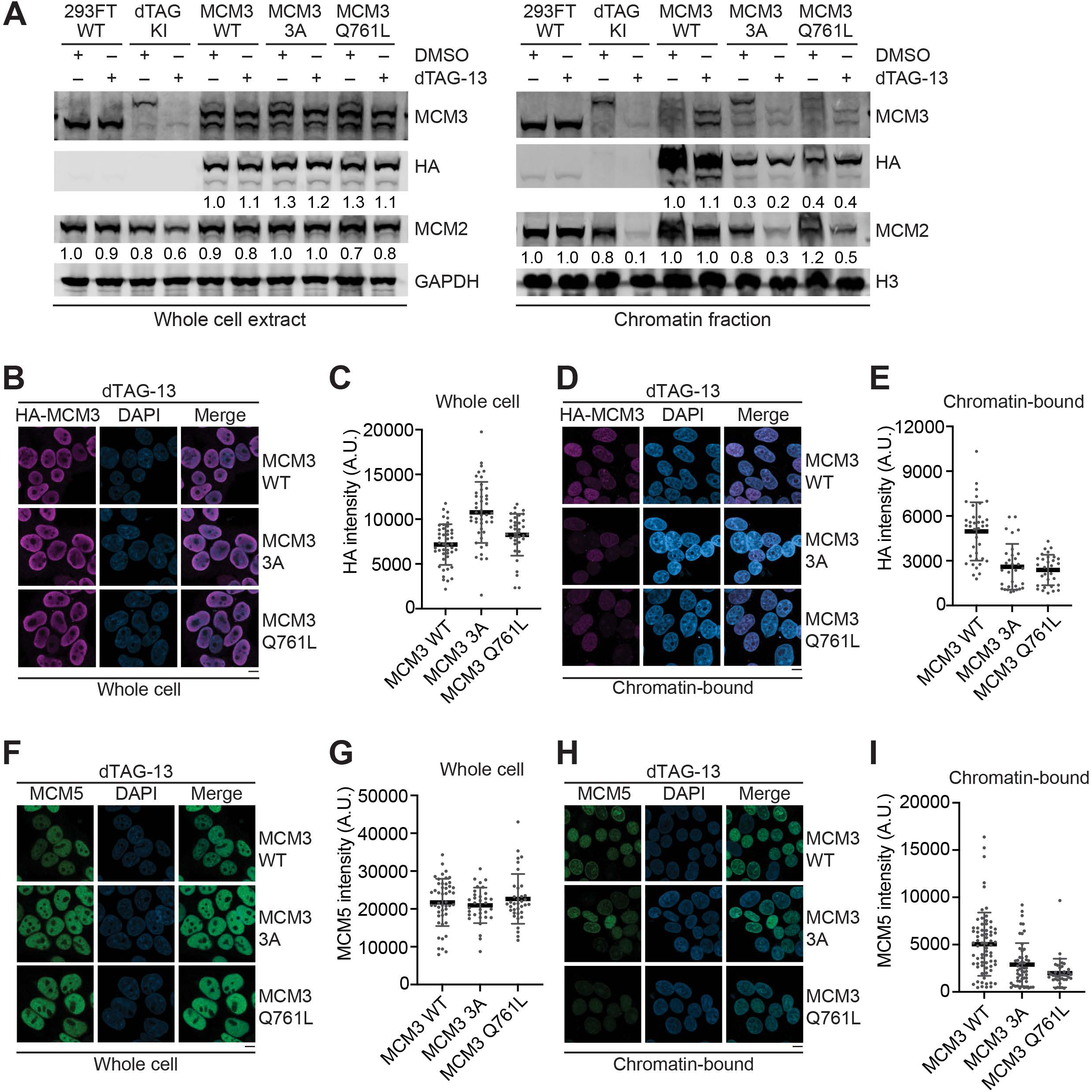
MCM3 WHD mutations reduce chromatin-bound MCM2–7 levels. (A) Immunoblots of whole-cell extracts (left panels) or chromatin fractions (right panels) of the indicated 293FT cells treated with DMSO or dTAG-13 (1 µM) for 16 h. GAPDH and histone H3 are used as loading controls. Relative intensities of HA-MCM3 and the endogenous MCM2 are quantified and shown below respective panels. (B) Images of indicated 293FT cells treated with dTAG-13 and stained with anti-HA (magenta) and DAPI (blue) without pre-extraction before fixation. The HA signals in this staining protocol represent total cellular HA-MCM3. (C) Quantification of HA-MCM3 intensities of cells in (B). Each dot in the graph represents a single cell. (D) Images of indicated 293FT cells treated with dTAG-13 and stained with anti-HA (magenta) and DAPI (blue) with pre-extraction before fixation. The HA signals in this staining protocol represent chromatin-bound HA-MCM3. (E) Quantification of HA-MCM3 intensities of cells in (D). (F) Images of indicated 293FT cells treated with dTAG-13 and stained with anti-MCM5 (green) and DAPI (blue) without pre-extraction before fixation. (G) Quantification of MCM5 intensities of cells in (F). (H) Images of indicated 293FT cells treated with dTAG-13 and stained with anti-MCM5 (green) and DAPI (blue) with pre-extraction before fixation. (I) Quantification of MCM5 intensities of cells in (H). For all relevant panels, the scale bar indicates 10 µm. Mean ± SD are shown.

In the yeast OCCM structure, Mcm3 WHD directly contacts the AAA+ ATPase domain of Cdc6.^10^ The interaction between Mcm3 WHD and Cdc6 contributes to Mcm2–7 loading. The reduced chromatin binding of MCM3 3A in human cells is likely due to the weakened interaction between MCM3 WHD and ORC-CDC6. Although the MCM3 Q761L mutation strengthens the interaction between MCM3 WHD and ORC-CDC6, it also strengthens the interaction between MCM3 WHD and MCM2 (i.e. the latch) that uses a similar interface. ORC-CDC6 may not be able to open the strengthened Q761L latch efficiently, resulting in an overall reduced interaction between ORC-CDC6 and the MCM2–7 complex containing the Q761L mutation.

We tested whether the residual loading of MCM3 3A and Q761L in human cells was still dependent on CDC6. CDC6 was efficiently depleted by transfection of a small interfering RNA against CDC6 (siCDC6) (Figure 5A). Consistent with results in Figure 4, without CDC6 depletion (siLUC transfection), chromatin-bound levels HA-MCM3 3A and Q761L were lower than that of WT in cells depleted of MCM3 (Figures 5B and 5C). CDC6 depletion reduced the chromatin-bound levels of HA-MCM3 WT, 3A, and Q761L to background levels, without affecting the total cellular levels of HA-MCM3 (Figures 5D and 5E). Thus, chromatin loading of MCM2–7 is strictly dependent on CDC6 in human cells. The deficient chromatin binding of MCM2–7 complexes containing MCM3 3A and Q761L still requires CDC6. Other molecular contacts between MCM2–7 and CDC6 that do not involve MCM3 WHD likely contribute to the loading of the MCM2–7 3A and Q761L mutants.

**Figure 5.**
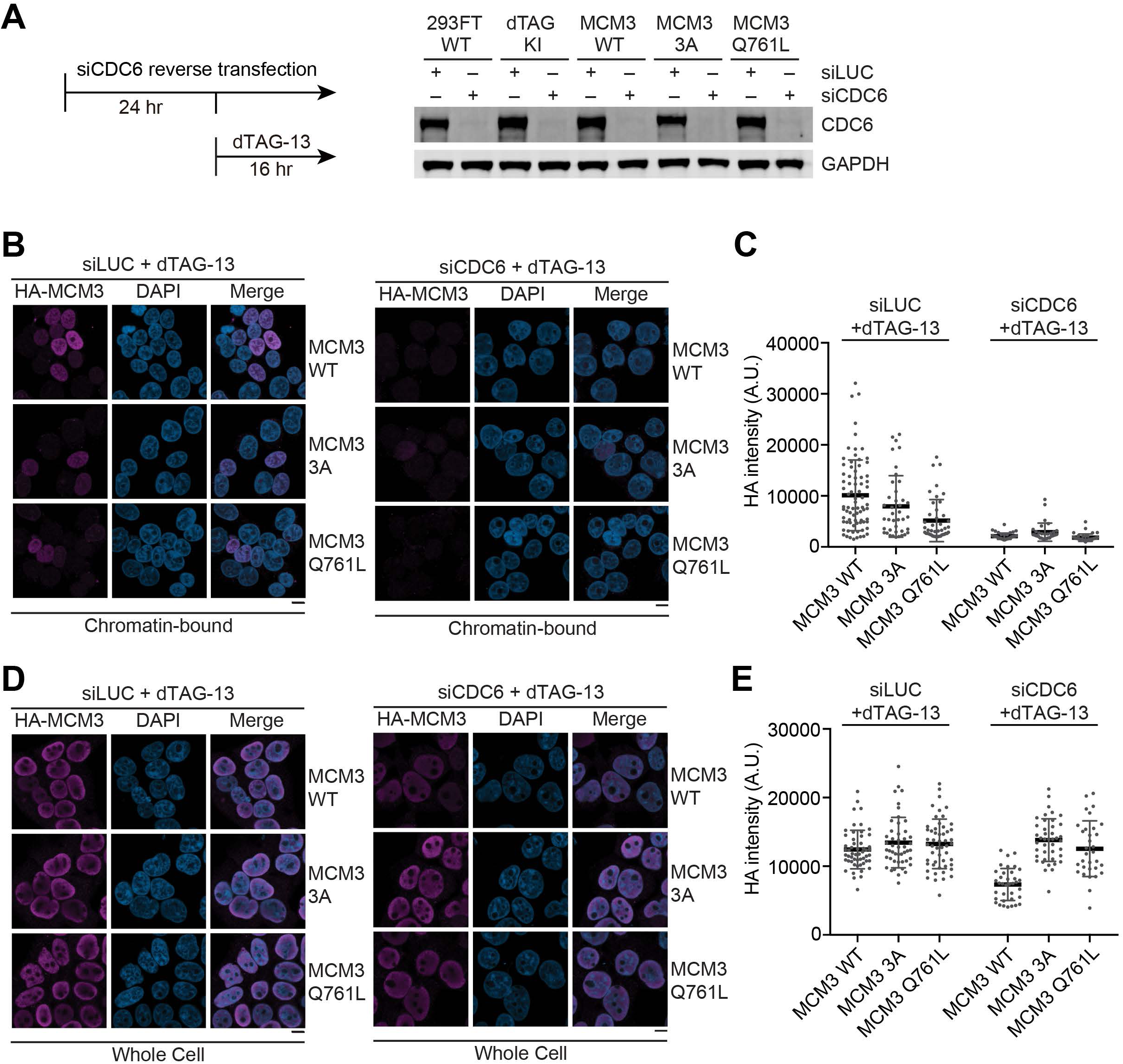
Loading of MCM2–7 harboring MCM3 WHD mutants depends on CDC6. (A) Immunoblots of whole-cell extracts of the indicated 293 FT cells that had been transfected with siLUC (as the negative control) or siCDC6 and then treated with dTAG-13 (1 µM) for 16 h. (B) Images of indicated 293FT cells transfected with siLUC (left panels) or siCDC6 (right panels), treated with dTAG-13, and then stained with anti-HA (magenta) and DAPI (blue) with pre-extraction before fixation. The HA signals in this staining protocol represent chromatin-bound HA-MCM3. (C) Quantification of HA-MCM3 intensities of cells in (B). Each dot in the graph represents a single cell. (D) Images of indicated 293FT cells transfected with siLUC (left panels) or siCDC6 (right panels), treated with dTAG-13, and then stained with anti-HA (magenta) and DAPI (blue) without pre-extraction before fixation. The HA signals in this staining protocol represent total cellular HA-MCM3. (E) Quantification of HA-MCM3 intensities of cells in (D). For all relevant panels, the scale bar indicates 10 µm. Mean ± SD are shown.

## Discussion

In this study, we determined the structures of human endogenous MCM2–7 complex with an open DNA entry gate between MCM2 and MCM5. We made two novel observations. First, we found that a fraction of human MCM2–7 complex in the absence of DNA formed a double hexamer (DH) with dynamic conformations. Whether the DNA-free MCM2–7 DH is functionally relevant for MCM2–7 loading remains to be tested in future experiments. Second, we showed that MCM3 WHD reaches across the DNA entry gate and binds to MCM2, thus creating a safety latch to prevent DNA from accessing the central channel. MCM3 WHD mutations that weaken or strengthen this latch cause deficient MCM2–7 loading and G2 arrest in human cells that is dependent on the DNA damage checkpoint.

We propose the following model to explain the complicated roles of MCM3 WHD in human MCM2–7 loading (Figure 6). DNA-unbound human MCM2–7 exists as both SH and DH with an open DNA entry gate between MCM2 and MCM5. In both the SH and DH, MCM3 WHD binds to MCM2, and together with the linker connecting WHD to the helicase domain, forming a safety latch to prevent DNA from accessing the central channel. DNA-bound ORC-CDC6 interacts with MCM3 WHD and recruits MCM2–7 to chromatin with the help of CDT1. ORC-CDC6 binding to MCM3 WHD simultaneously opens the safety latch, allowing DNA to access the central channel. DNA binding also displaces MCM4 and MCM5 WHDs from the central channel. The DNA entry gate of MCM2–7 then closes to entrap DNA, forming the pre-replicative OCCM complex.

**Figure 6.**
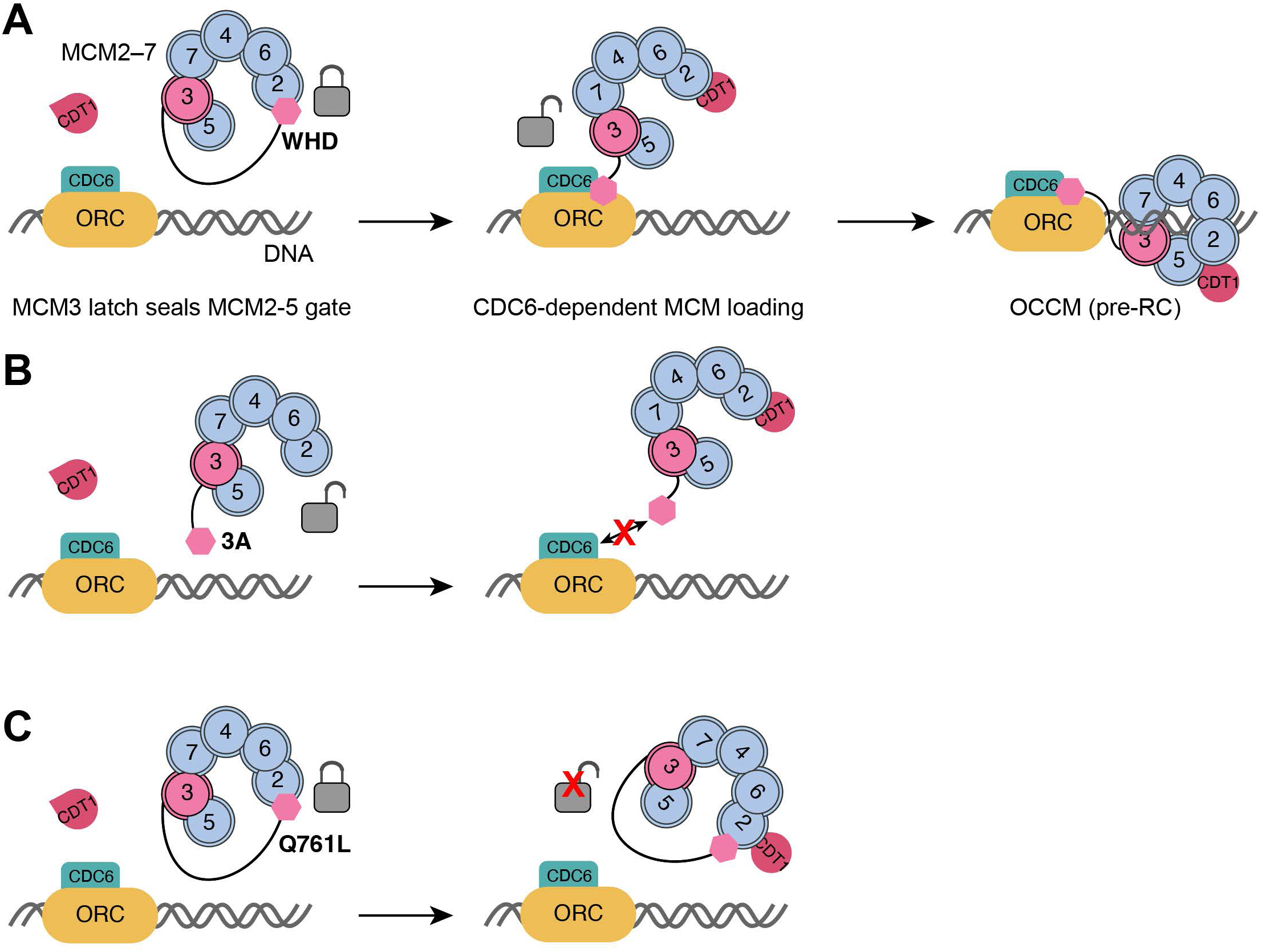
Model explaining the functions of MCM3 WHD. (A) In the pre-loading state of MCM2–7, MCM3 WHD functions as a latch to block DNA entry through the MCM2-5 gate. Binding of MCM3 WHD to the ORC-CDC6 complex recruits MCM2–7 to chromatin and simultaneously opens the WHD latch, allowing DNA entry into the MCM2–7 central channel and the formation of the ORC-CDC6-CDT1-MCM2–7 (OCCM) complex on chromatin. (B) MCM3 WHD 3A mutations disrupt the binding of MCM3 WHD to MCM2 and open the latch. MCM2–7 with the opened latch is, however, not prematurely loaded onto DNA. Because MCM3 WHD 3A is deficient in binding to ORC-CDC6, MCM2–7 carrying the 3A mutation is in fact less efficiently loaded onto DNA, causing replication defects in human cells. (C) The disease-related MCM3 WHD Q761L mutation strengthens the latch, which cannot be properly opened by ORC-CDC6. This mutation also causes MCM2–7 loading defects and DNA replication stress in cells.

MCM2–7 containing the MCM3 WHD^3A^ mutant, which is deficient for MCM2 binding, has an opened latch, but does not exhibit CDC6-independent loading on chromatin in human cells. Thus, latch opening alone is not sufficient to cause spontaneous loading of MCM2–7 onto DNA. Other conformational changes mediated by ORC-CDC6 are still required to load MCM2–7. In fact, because MCM3 WHD^3A^ has a weaker affinity towards ORC-CDC6, MCM2–7 with the open latch is loaded less efficiently on chromatin (Figure 6B). Interestingly, the MCM3 WHD^Q761L^ has been linked to a human developmental disease called Meier-Gorlin syndrome.^25^ This mutation increases the hydrophobicity of the MCM3 WHD-MCM2 interface and the MCM3 WHD-ORC-CDC6 interface, thus strengthening both interactions. The net outcome is that the strengthened WHD^Q761L^ latch cannot be efficiently opened by ORC-CDC6 (Figure 6C), leading to MCM2–7 loading defects. Thus, the MCM3 WHD latch, when locked, prevents DNA entry into the MCM2–7 central channel.

Both MCM3 WHD^3A^ and WHD^Q761L^ mutations elicit G2 arrest in human cells that is dependent on the ATR-CHK1 branch of the DNA damage checkpoint. This finding suggests that these mutations might cause replication stress and incomplete DNA replication during S phase. Because MCM2–7 complexes are loaded in excess onto chromatin before S phase in human cells, not all MCM2–7 complexes are activated to become CMG helicases and generate replication forks.^31–34^ It is thus unclear whether the relatively mild loading defects of MCM2–7 complexes carrying MCM3 WHD^3A^ and WHD^Q761L^ mutations suffice to explain the strong cell cycle defects. This MCM3 WHD interface might have additional functions during DNA replication.

In conclusion, we have identified the opening of the MCM3 WHD safety latch as a previously uncharacterized step needed for human MCM2–7 loading and demonstrated its functional relevance in human cells. Future experiments are needed to test whether the MCM2– 7 DH in the pre-loading state is a functionally relevant loading intermediate during pre-RC formation.

## Methods

### Key Resources Table

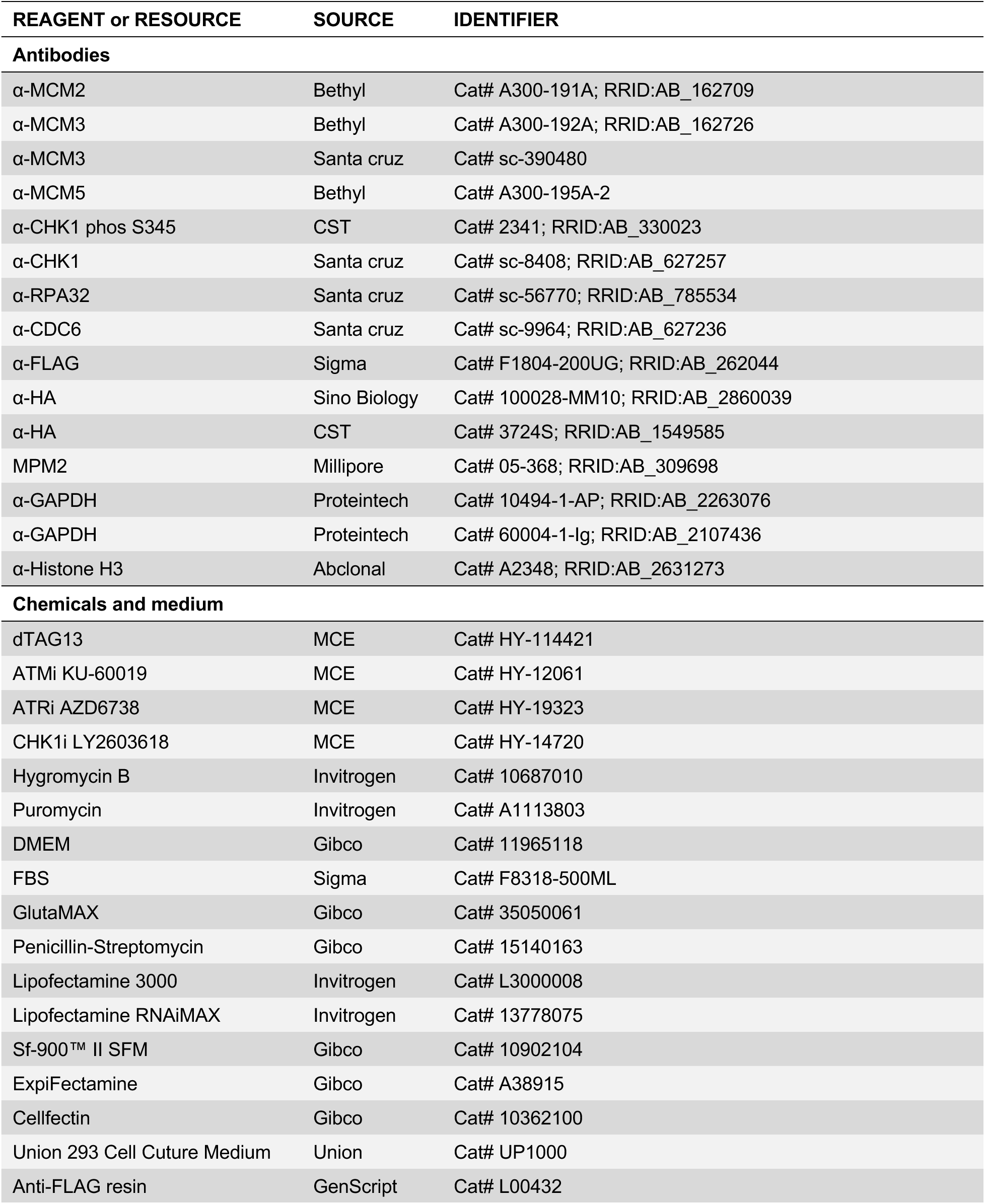
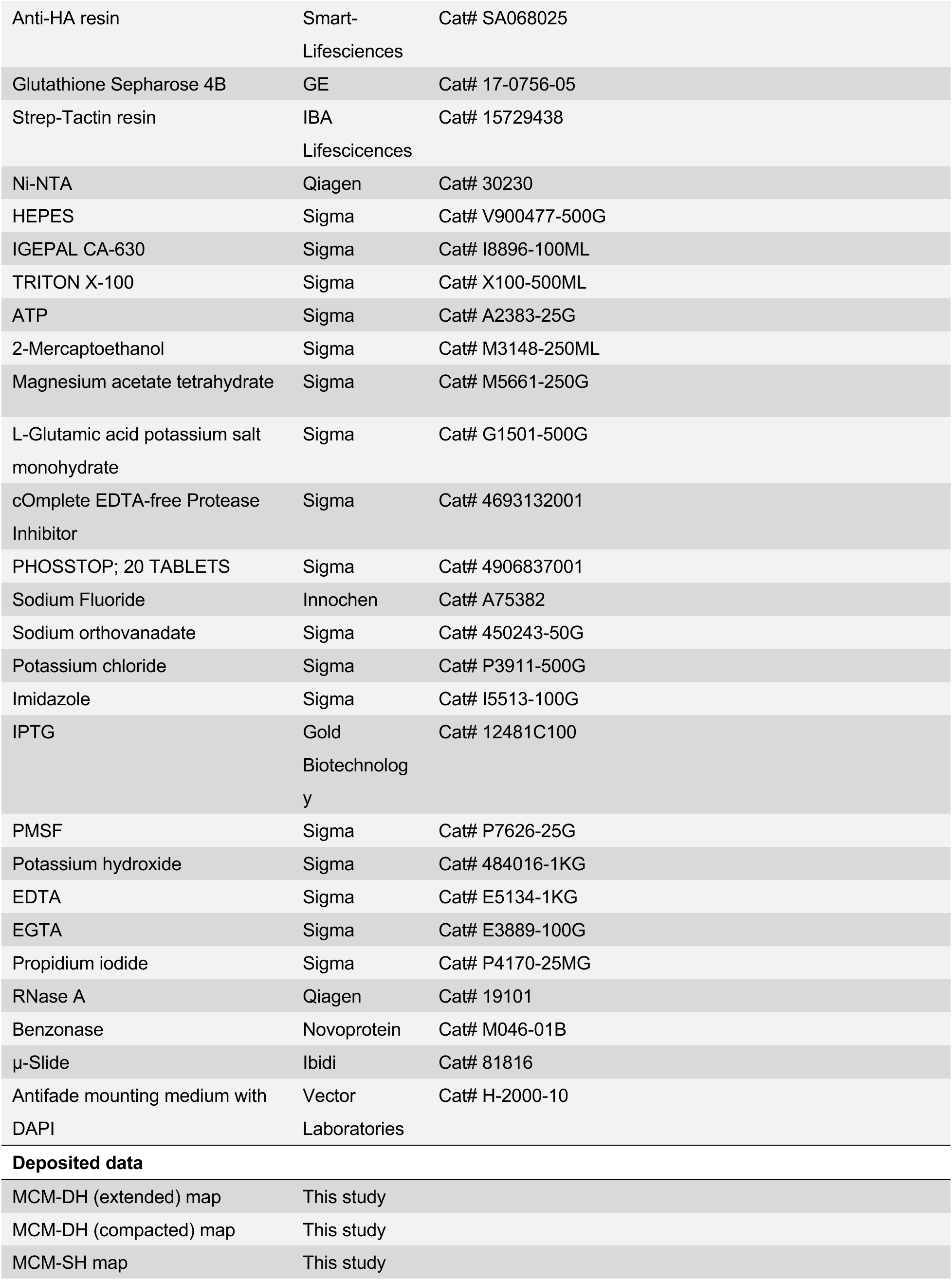
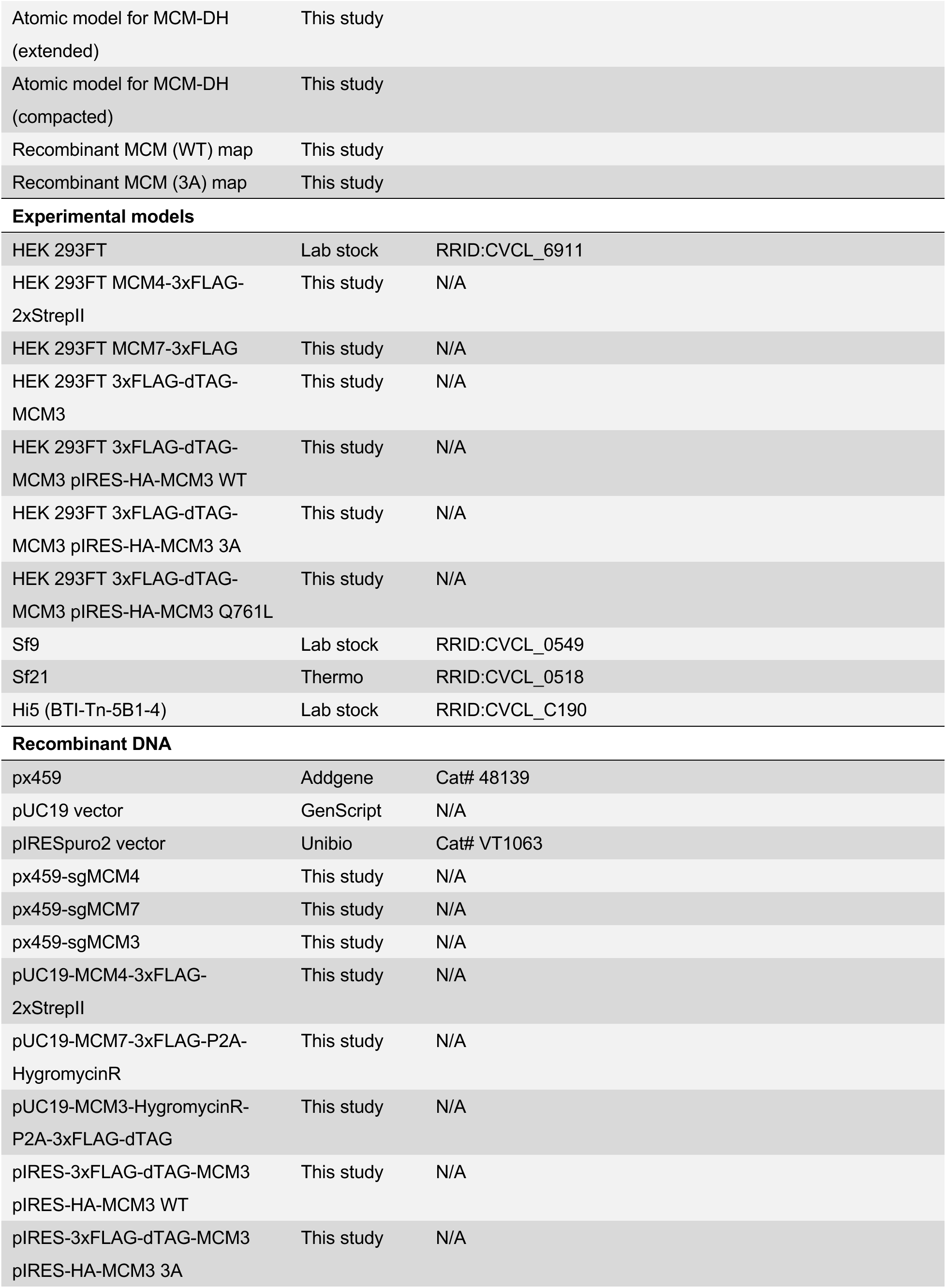
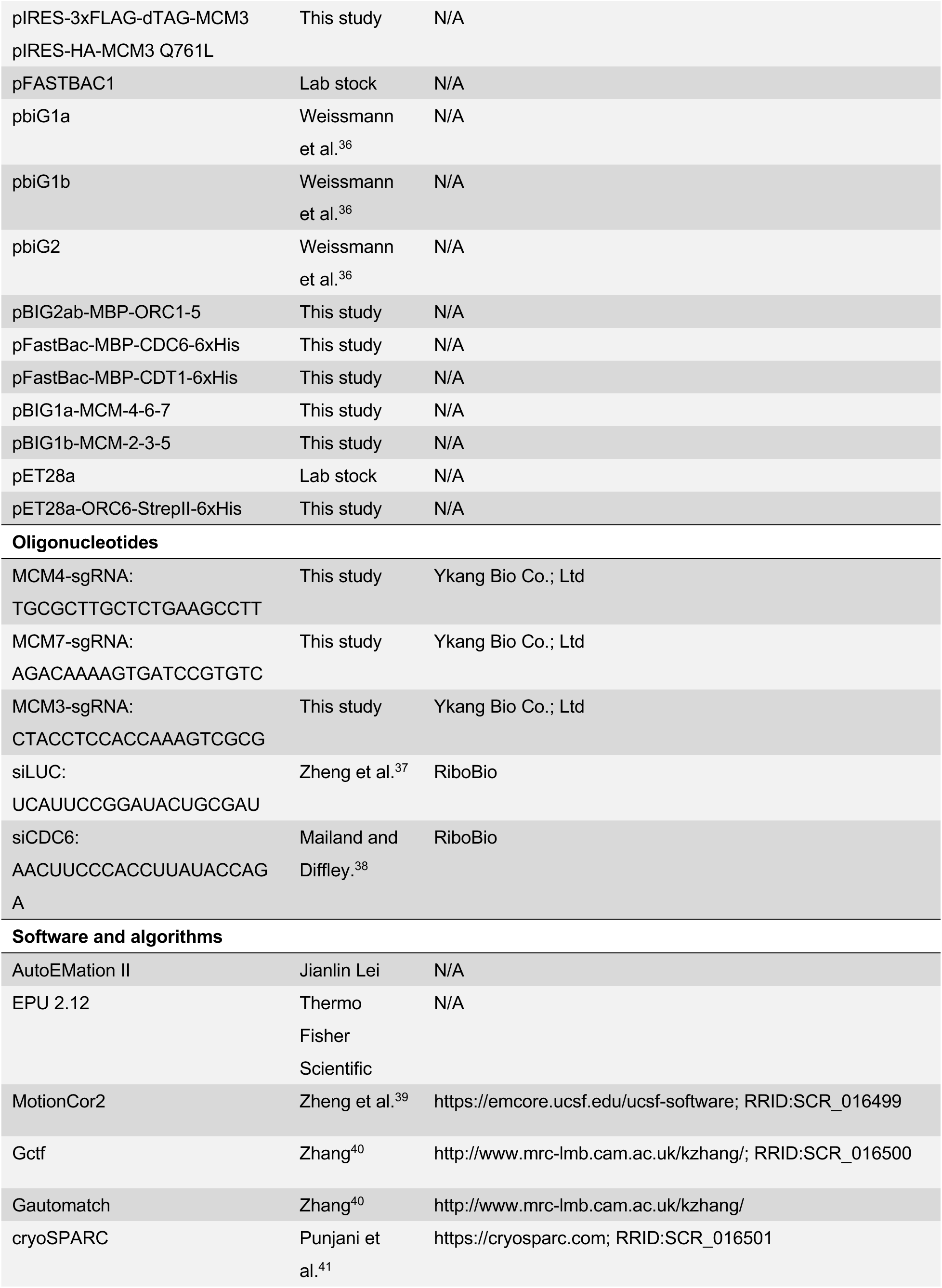
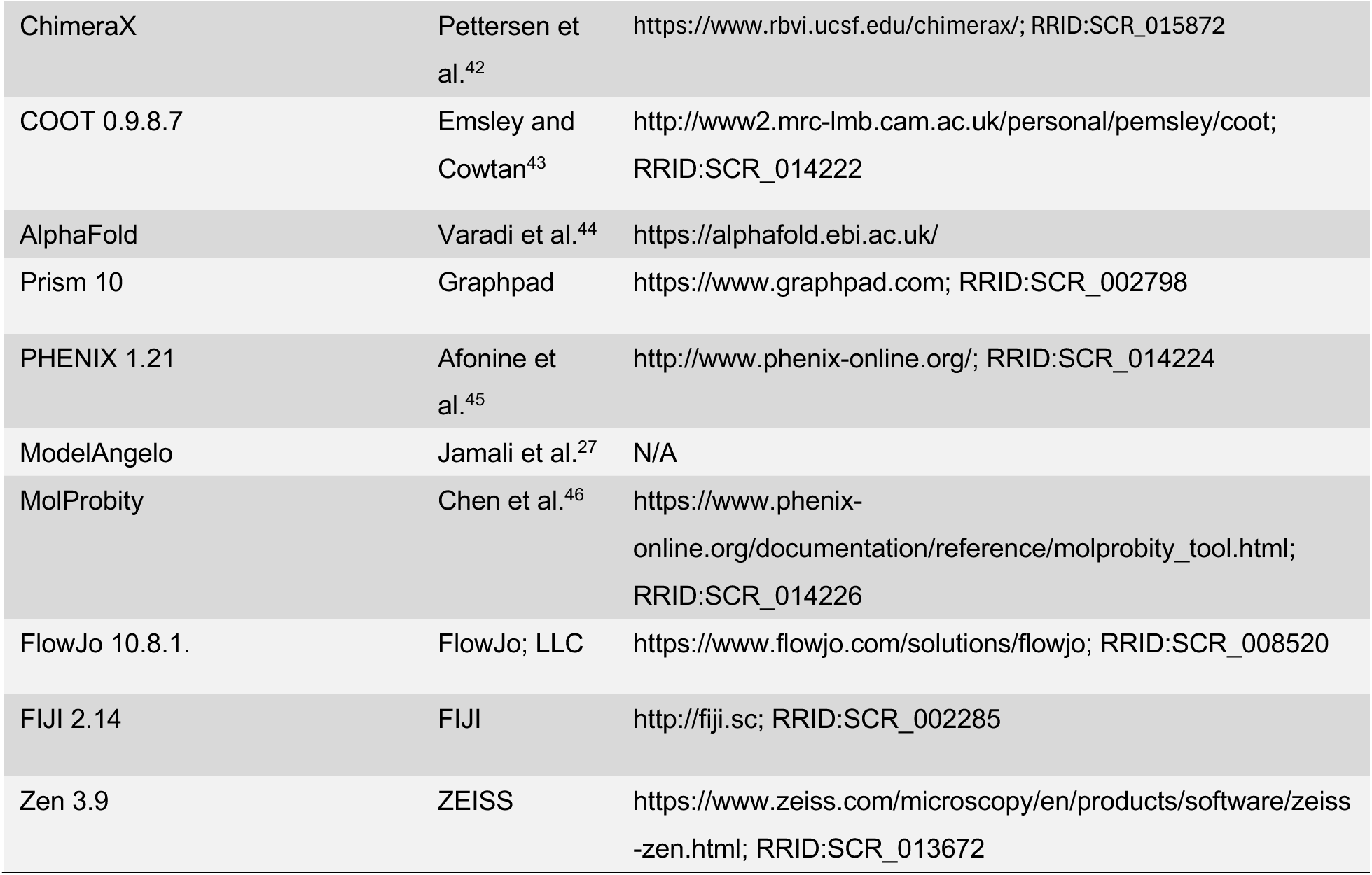

### Cell culture and cell lines

Human 293FT cells were cultured in the DMEM medium supplemented with 10% fetal bovine serum (FBS) and cultured at 37°C with 5% CO_2_. MCM4-3xFLAG-2xStrepII, MCM7-3xFLAG and 3xFLAG-dTAG-MCM3 293FT cells were generated in this study by using the CRISPR/Cas9 system. The correct tagging was confirmed by immunoblotting and/or DNA sequencing. 3xFLAG-dTAG-MCM3 293FT cells stably expressing HA-MCM3 WT, HA-MCM3 3A, and HA-MCM3 Q761L were constructed by transfecting 3xFLAG-dTAG-MCM3 293FT cells with mammalian expression plasmids and selected by the appropriate antibiotics.

For creating MCM4-3xFLAG-2xStrepII 293FT cells, the MCM4 sgRNA (TGCGCTTGCTCTGAAGCCTT) was ligated into the PX459 vector (Addgene; 48139). The donor sequence of 3xFLAG-2xStrepII flanked by homology arms of the *MCM4* genomic sequence was synthesized and cloned into the pUC19 vector. The PX459-sgMCM4 and pUC19-MCM4-3xFLAG-2xStrepII plasmids were co-transfected into 293FT cells using Lipofectamine 3000 (Invetrogen; L3000008) according to the manufacturer’s instructions. Cells were selected with 1 ug/ml puromycin (Invitrogen; A1113803) for 7 days. For making MCM7-3xFLAG 293FT cells, the MCM7 sgRNA (AGACAAAAGTGATCCGTGTC) was ligated into the PX459 vector. The donor sequence of 3xFLAG-P2A-HygromycinR flanked by homology arms of the *MCM7* genomic sequence was synthesized and cloned into the pUC19 vector. The PX459-sgMCM7 and pUC19-MCM7-3xFLAG-P2A-HygromycinR plasmids were co-transfected into 293FT cells. Cells were selected with 200 ug/ml hygromycin B (Invitrogen; 10687010) for 7 days. For making 3xFLAG-dTAG-MCM3 293FT cells, the MCM3 sgRNA (CTACCTCCACCAAAGTCGCG) was ligated into PX459 vector. The donor sequence of HygromycinR-P2A-3xFLAG-dTAG flanked by the homology arms of the *MCM3* genomic sequence was synthesized and cloned into pUC19. The PX459-sgMCM3 and pUC19-MCM3-HygromycinR-P2A-3xFLAG-dTAG plasmids were co-transfected into 293FT cells. Cells were selected with 200 ug/ml hygromycin B for 7 days. The surviving cells were sorted into 96-well plates. Clones with the correct targeting were validated by PCR and DNA sequencing.

For generating 3xFLAG-dTAG-MCM3 293FT cells stably expressing HA-MCM3 WT, HA-MCM3 I804A/F806A/I808A (3A), or HA-MCM3 Q761L, the 3A and Q761L mutations were introduced into the pIRESpuro2-3xHA-MCM3 vector with the Q5 Site-Directed Mutagenesis Kit (NEB; E0554S). The plasmids were transfected into 3xFLAG-dTAG-MCM3 293FT cells using Lipofectamine 3000. Cells were selected with 1 ug/ml puromycin (Invitrogen; A1113803) for 7 days. Transgene expression was validated by immunoblotting. The stable cell pools were maintained with media containing 0.7 ug/ml puromycin for future experiments.

For depletion of endogenous MCM3, dTAG-13 was added to the medium at a final concentration of 1 µM and incubated for an additional 16 hours, with DMSO treatment included as the vehicle control. For CDC6 depletion, cells were reverse transfected using Lipofectamine RNAiMAX with 50 nM siCDC6 (AACUUCCCACCUUAUACCAGA) or siLUC (UCAUUCCGGAUACUGCGAU) while seeding. After 3 hours, dTAG-13 was added to the medium at a final concentration of 1 µM and incubated for an additional 16 hours. For checkpoint inhibitor treatment, cells were treated with DMSO (as the vehicle control), 200 nM ATMi (KU-60019), ATRi (AZD6738), or CHK1i (LY2603618).

### Protein expression and purification

The endogenous human MCM2–7 complex from 293FT cells was purified as previously described with minor modifications.^24^ In brief, 293FT cells with affinity-tagged MCM4 or MCM7 were harvested by centrifugation at 3,990 g for 5 min. Cell pellets were resuspended and lysed by the extraction buffer [50 mM HEPES pH 7.5, 10 mM Mg(OAc)_2_, 100 mM potassium gluconate (K-Glu), 3 mM ATP, 1 mM EDTA. 0.5% Triton X-100, 2 mM NaF, 1 mM Na_3_VO_4_, 1 mM PMSF, 1x cOmplete EDTA-free Protease Inhibitor Cocktail]. Chromatin was collected and treated with 1U/µl benzonase (Novoprotein) in the extraction buffer supplemented with 8 mM MgCl_2_. The soluble fraction was then incubated with anti-FLAG beads (GenScript). After wash, the MCM2– 7 complexes bound to beads were released by Precision Protease cleavage, concentrated, and fractionated by a 20%-40% glycerol gradient (for the MCM4-3xFLAG-2xStrepII complex) or a 15%-50% sucrose gradient (for the MCM7-3xFLAG complex) in the extraction buffer with 1x cOmplete EDTA-free Protease Inhibitor Cocktail in an SW-60Ti rotor (Beckman Optima XPN-100 Ultracentrifuge) at 126,100 g for 14 h. The peak fractions were concentrated, aliquoted, and stored at −80_°_C.

For the expression and purification of recombinant human ORC1-6 complex, full-length, codon-optimized synthetic cDNAs of human ORC1-6 subunits were cloned into the BIGBac baculovirus expression system. Multiple tags were added to different subunits: an MBP-tag, twin-StrepII, and SumoStar tag at the N-terminus of ORC1; a 3xFlag-tag at the N-terminus of ORC2; and HA-tags at the C-terminus of ORC3 and ORC5. Sf21 cells infected by the ORC1-6 baculovirus were collected and lysed in the high-salt buffer [25 mM HEPES, pH 7.6, 0.8 M KCl, 5 mM ATP, 10% glycerol, 0.05% (v/v) IGEPAL CA-630, 1×Protease inhibitor cocktail] and resuspended in the low-salt buffer [25 mM HEPES, pH 7.6, 0.4 M KCl, 5 mM ATP, 10% glycerol, 0.05% (v/v) IGEPAL CA-630, 1×cOmplete EDTA-free Protease Inhibitor Cocktail]. The ORC complex was purified with tandem affinity purification using Streptactin Beads (Smart-Lifesciences) and anti-HA beads (Smart-Lifesciences), released by TEV protease cleavage, and fractionated by size exclusion chromatography (Superdex200, GE). Peak fractions were concentrated, aliquoted, and stored at −80°C.

ORC6 tagged with 2xStrepII and His_6_ tag was expressed in E. coli upon induction with 0.5 mM Isopropyl β-D-1-thiogalactopyranoside (IPTG) overnight at 18°C. The bacteria were harvested by centrifugation and resuspended in the ORC6 lysis buffer [50 mM Tris-HCl, pH 7.0, 300 mM NaCl, 10% glycerol, 0.05% (v/v) IGEPAL CA-630, and 1×Protease inhibitor cocktail]. After sonication and centrifugation, the supernatant was incubated with Streptactin Beads and eluted with the ORC6 lysis buffer containing 5 mM biotin. The eluate was fractionated by cation exchange chromatography (Resource S, GE) and gel filtration (Superdex75, GE). Peak fractions were concentrated, aliquoted, and stored at −80°C.

Recombinant human CDC6 and CDT1 proteins were individually expressed and purified from insect cells. Full-length CDC6 and CDT1 were tagged with MBP and the Prescission protease cleavage site at the N-terminus, and His_6_ at the C-terminus. Cells were collected and lysed with the lysis buffer (50 mM HEPES-KOH, pH 7.8, 600 mM KCl, 10% glycerol, 1 mM DTT, 25 mM imidazole). CDC6 and CDT1 proteins were tandem-affinity purified by Ni-NTA beads (Qiagen) and amylose affinity beads (NEB). The eluate was further fractionated by gel filtration (Superdex200, GE). Proteins were concentrated, aliquoted, and stored at −80°C.

For expression of recombinant will-type and mutant human MCM2–7 complexes, MCM2– 7 subunits were cloned into two biGBac expression vectors: one containing MCM3, MCM5, and MCM7, and the other containingMCM2, MCM4 (with a prescission protease cleavage site and 3xFLAG tag at the C-terminus), and MCM6. Hi5 insect cells were infected with these baculoviruses, harvested, lysed in the MCM lysis buffer (50 mM HEPES-KOH, pH 7.5, 100 mM potassium glutamate, 10% glycerol, pH 7.5, 10 mM MgCl2, 1 mM 2-Mercaptoethanol, 200 mM PMSF, 5 mM ATP). After sonication and centrifugation, the supernatant was incubated with anti-FLAG beads. MCM2–7 was released from washed beads by Prescission Protease cleavage. The eluate was fractionated by gel filtration (Superose6, GE). Peak fractions were concentrated, aliquoted, and stored at −80°C.

### Antibodies

The following antibodies against human proteins were used for immunoblotting and immunofluorescence: MCM2 (Bethyl Laboratories; A300-191A), MCM3 (Bethyl Laboratories; A300-192A), MCM3 (Santa Cruz; sc-390480), MCM5 (Bethyl Laboratories; A300-195A-2), CHK1 phospho-S345 (CST; 2341), CHK1 (Santa Cruz; sc-8408), RPA32 (Santa Cruz; sc-56770), CDC6 (Santa Cruz; sc-9964), FLAG (Sigma; F1804-200UG), HA (Sino Biology; 100028-MM10), HA (CST; 3724S), GAPDH (Proteintech; 10494-1-AP and 60004-1-Ig), Histone H3 (Abclonal; A2348).

### Immunoblotting

After 24 hours, dTAG-13 was added to the medium at a final concentration of 1 µM and incubated for an additional 16 hours, with DMSO as the vehicle control. For preparation of whole cell lysates, cell pellets were collected, lysed in the Laemmli sample buffer, and denatured at 95°C. Chromatin fractionation was performed as described previously.^47^ In brief, the cells were suspended at a concentration of 4 × 10^7^ cells/ml in buffer A (10 mM HEPES-KOH pH 7.9, 10 mM KCl, 1.5 mM MgCl_2_, 0.34 M sucrose, 10% glycerol, 1 mM DTT, 1 x PhosphoSTOP, and 1 x cOmplete EDTA-free Protease Inhibitor Cocktail) with 0.1% Triton X-100 and incubated on ice for 5 min. The nuclei were pelleted with low-speed centrifugation at 1,300 g for 4 min at 4°C, washed once with buffer A and twice with buffer B (3 mM EDTA, 0.2 mM EGTA, 1 mM DTT, 1 x PhosphoSTOP, and 1 x cOmplete EDTA-free Protease Inhibitor Cocktail) via centrifugation at 1,700 g for 4 min at 4°C. The chromatin fraction was lysed in the Laemmli sample buffer and denatured at 95°C. Both whole cell lysates and chromatin fractions were separated by SDS-PAGE and blotted with the desired primary antibodies. Anti-mouse IgG (H+L) (DyLight 800 4X PEG Conjugate) (CST; 5257) and anti-rabbit IgG (H+L) (DyLight 680 Conjugate) (CST; 5366) were used as secondary antibodies. The blots were scanned with an Odyssey Infrared Imaging System (LI-COR).

### Immunofluorescence

Cells were cultured and treated in the µ-Slide 18 Well Chamber Slides (ibidi). Cells were washed once with PBS and then fixed in 4% paraformaldehyde for 15 min. For staining chromosome-bound proteins, cells on the chamber slides were washed once with PBS, pre-extracted with ice-cold PBS supplemented with 0.5% Triton X-100 for 5 min, and then fixed in 4% paraformaldehyde for 15 min. The fixed cells were blocked in PBS with 0.1% Triton X-100 (PBST) containing 3% BSA at room temperature for 30 min and incubated with the desired antibodies in PBST containing 3% BSA at room temperature for 2 h. After three washes with PBST, cells were incubated with fluorescent secondary antibodies in PBST containing 3% BSA for 1 h at room temperature and washed three times with PBST. After the final wash of PBST, the slides were mounted with VECTASHIELD PLUS Antifade Mounting Medium with DAPI (H-2000) and viewed with a 63X objective on a ZEISS LSM 900 with Airyscan 2 (ZEISS). Image processing and quantification were performed with Zen 3.9 (ZEISS) and ImageJ (adjusted by FIJI).

### Flow cytometry

Cells were collected by trypsinization and fixed in 70% ice-cold ethanol overnight. After fixation, cells were washed once with PBST, resuspended in PBST supplemented with 20 µg/ml propidium iodide (Sigma) and 200 µg/ml RNase A (Qiagen), and incubated at room temperature for 1 h. The samples were analyzed on a CytoFLEX LX-6L flow cytometer (Beckman). Data processing and quantification were performed with CytExpert and FlowJo 10.8.1. For MPM2 staining, fixed cells were washed once with PBST and incubated with the MPM2 antibody (1:200, Millipore) in PBST containing 1% BSA for 3 h. Cells were then washed once with PBST and incubated with the fluorescent secondary antibody in PBST containing 1% BSA for 30 min, followed by propidium iodide staining and analysis on a flow cytometer. The percentage of cells that had 4N DNA content and were MPM2-positive was defined as the mitotic index.

### Cryo-EM data collection and image processing

For cryo-EM grid preparation, 3 µl samples (∼5 mg/ml) were applied onto glow-discharged holey carbon grids (Quantifoil Cu R1.2/1.3, 300 mesh), blotted with a Vitrobot Marker IV (Thermo Fisher Scientific) for 3 s under 100% humidity at 4°C, and subjected to plunge freezing into liquid ethane. All cryo-EM data were collected using an FEI Titan Krios microscope at 300 kV equipped with a Gatan K3 Summit direct electron detector (in super-resolution mode and at a nominal magnification of 81,000) and a GIF-quantum energy filter. Defocus values were set from -1.0 to -2.0 um. Each stack of 32 frames was exposed for 2.56 s, with a total electron dose of 50 e^-^/Å^2^. AutoEMation was used for fully automated data collection.

All micrograph stacks were motion corrected with MotionCor2 with a binning factor of 2, resulting in a pixel size of 1.0773 Å or 1.087 Å, respectively. Contrast transfer function (CTF) parameters were estimated using Gctf.^40^ Most steps of image processing were performed using cryoSPARC.^41^ For 3D processing of the data, particles were automatically picked from micrographs using Gautomatch (developed by Kai Zhang, MRC-LMB, http://www.mrc-lmb.cam.ac.uk/kzhang/). Particles were extracted with a pixel size of 4.3092 Å and subjected to several rounds of reference-free 2D classification. Particles were kept after the exclusion of obvious ice contamination and junk particles. The particles were reextracted without binning. Reference-free 2D classification was performed to exclude the junk particles.

For the extended conformation, ab-initio models were generated and used for heterogeneous 3D refinement. The best class of 133,948 particles was then used for further homogenous non-uniform refinement, and local refinement with C2 symmetry. The global resolution is 3.86 Å based on the Fourier Shell Correlation (FSC) 0.143 criterion. For the compacted conformation, heterogenous refinement was performed with imported initial models. The best class of 254,512 particles was then used for further non-uniform and local refinement, followed by further 2D classification. Finally, 242,055 particles were used for the non-uniform and local refinement with C2 symmetry. The global resolution is 3.96 Å based on the Fourier Shell Correlation (FSC) 0.143 criterion. Heterogeneous and non-uniform refinements were also performed without applying symmetry, and local refinement masked for one of the single hexamers was performed to generate a higher resolution single hexamer map with a global resolution at 3.27 Å based on the Fourier Shell Correlation (FSC) 0.143 criterion.

### Model building and refinement

AlphaFold3 was used to generate the starting model and docked into the final EM maps with UCSF ChimeraX.^42^ The models were manually adjusted and iteratively built in COOT^43^ and then refined against summed maps using phenix.real_space_refine implemented in PHENIX^45^ until the validation data were reasonable. FSC values were calculated between the resulting models and the two half-maps, as well as the averaged map of the two half-maps. The quality of the models was evaluated with MolProbity.^46^ The structure validation statistics were listed in Table S1. All structural figures were prepared with ChimeraX.^42^ Map-to-model autobuilding was performed with ModelAngelo.^27^

### Statistical analysis

No statistical methods were used to predetermine sample size or applied to data analysis. The experiments were not randomized. The investigators were not blinded to allocation during experiments and outcome assessment.

## Data availability

The cryo-EM density maps of the extended MCM2–7 DH, compacted MCM2–7 DH, masked MCM2–7 SH, recombinant MCM2–7, and recombinant MCM2–7 containing the MCM3 3A mutant have been deposited to the Electron Microscopy Data Bank under the accession numbers EMD-63476 (conformer I; extended), EMD-63475 (conformer II; compacted), EMD-63474 (SH), EMD-63486 (recombinant WT), and EMD-63487 (recombinant 3A). Atomic coordinates have been deposited to the RCSB Protein Data Bank under the accession numbers 9LXF (extended DH), 9LXE (compacted DH), and 9LXD (SH).

## Acknowledgements

Cryo-EM data were collected at the Westlake University Cryo-EM Facility. We thank Jianlin Lei, Zhipeng Jiang, Li Huang, Yinling Zhang, Xiaojuan Wang, and Yali Gu for technical support and facility access. We thank Biomedical Research Core Facilities for technical support on flow cytometry. We thank the Westlake University High-Performance Computing Center for computational resources and technical assistance. This work was supported by the National Natural Science Foundation of China (Project 32130053 to H.Y. and Project 32271258 to H.G.) the “Pioneer” and “Leading Goose” R&D Program of Zhejiang (2024SSYS0036 to H.Y.), and the Tencent New Cornerstone Science Foundation (to H.Y.).

## Author contributions

Y.L. and M.Y. purified proteins and performed biochemical assays. Y.L., M.Y., P.L., and H.G. performed cryo-EM grid preparation, data collection, and processing. Y.L. and H.G. performed model building. Y.L. performed all cellular assays. Y.L. and all authors analyzed the data. H.Y. and Y.S. designed and supervised the project. Y.L., H.G., and H.Y. wrote the manuscript with input from all authors.

## Competing interests

The authors declare no competing interests.

## Notes

### Competing Interest Statement

The authors have declared no competing interest.

## References

1. Hu, Y., and Stillman, B. (2023). Origins of DNA replication in eukaryotes. Mol Cell 83, 352–372. 10.1016/j.molcel.2022.12.024.

2. Costa, A., and Diffley, J.F.X. (2022). The Initiation of Eukaryotic DNA Replication. Annu Rev Biochem 91, 107–131. 10.1146/annurev-biochem-072321-110228.

3. Tye, B.K., and Zhai, Y. (2023). The Origin Recognition Complex: From Origin Selection to Replication Licensing in Yeast and Humans. Biology (Basel) 13. 10.3390/biology13010013.

4. Lewis, J.S., Gross, M.H., Sousa, J., Henrikus, S.S., Greiwe, J.F., Nans, A., Diffley, J.F.X., and Costa, A. (2022). Mechanism of replication origin melting nucleated by CMG helicase assembly. Nature 606, 1007–1014. 10.1038/s41586-022-04829-4.

5. Zhai, Y., Li, N., Jiang, H., Huang, X., Gao, N., and Tye, B.K. (2017). Unique Roles of the Non-identical MCM Subunits in DNA Replication Licensing. Mol Cell 67, 168–179. 10.1016/j.molcel.2017.06.016.

6. Li, N., Zhai, Y., Zhang, Y., Li, W., Yang, M., Lei, J., Tye, B.K., and Gao, N. (2015). Structure of the eukaryotic MCM complex at 3.8 A. Nature 524, 186–191. 10.1038/nature14685.

7. Zhai, Y., Cheng, E., Wu, H., Li, N., Yung, P.Y., Gao, N., and Tye, B.K. (2017). Open-ringed structure of the Cdt1-Mcm2-7 complex as a precursor of the MCM double hexamer. Nat Struct Mol Biol 24, 300–308. 10.1038/nsmb.3374.

8. Yuan, Z., Schneider, S., Dodd, T., Riera, A., Bai, L., Yan, C., Magdalou, I., Ivanov, I., Stillman, B., Li, H., and Speck, C. (2020). Structural mechanism of helicase loading onto replication origin DNA by ORC-Cdc6. Proc Natl Acad Sci U S A 117, 17747–17756. 10.1073/pnas.2006231117.

9. Xu, N., Lin, Q., Tian, H., Liu, C., Wang, P., Suen, C.M., Yang, H., Xiang, Y., and Zhu, G. (2022). Cryo-EM structure of human hexameric MCM2-7 complex. iScience 25, 104976. 10.1016/j.isci.2022.104976.

10. Yuan, Z., Riera, A., Bai, L., Sun, J., Nandi, S., Spanos, C., Chen, Z.A., Barbon, M., Rappsilber, J., Stillman, B., et al. (2017). Structural basis of Mcm2-7 replicative helicase loading by ORC-Cdc6 and Cdt1. Nat Struct Mol Biol 24, 316–324. 10.1038/nsmb.3372.

11. Faull, S.V., Barbon, M., Mossler, A., Yuan, Z., Bai, L., Reuter, L.M., Riera, A., Winkler, C., Magdalou, I., Peach, M., et al. (2025). MCM2-7 ring closure involves the Mcm5 C-terminus and triggers Mcm4 ATP hydrolysis. Nat Commun 16, 14. 10.1038/s41467-024-55479-1.

12. Feng, X., Noguchi, Y., Barbon, M., Stillman, B., Speck, C., and Li, H. (2021). The structure of ORC-Cdc6 on an origin DNA reveals the mechanism of ORC activation by the replication initiator Cdc6. Nat Commun 12, 3883. 10.1038/s41467-021-24199-1.

13. Schmidt, J.M., and Bleichert, F. (2020). Structural mechanism for replication origin binding and remodeling by a metazoan origin recognition complex and its co-loader Cdc6. Nat Commun 11, 4263. 10.1038/s41467-020-18067-7.

14. Schmidt, J.M., Yang, R., Kumar, A., Hunker, O., Seebacher, J., and Bleichert, F. (2022). A mechanism of origin licensing control through autoinhibition of S. cerevisiae ORC.DNA.Cdc6. Nat Commun 13, 1059. 10.1038/s41467-022-28695-w.

15. Sun, J., Evrin, C., Samel, S.A., Fernandez-Cid, A., Riera, A., Kawakami, H., Stillman, B., Speck, C., and Li, H. (2013). Cryo-EM structure of a helicase loading intermediate containing ORC-Cdc6-Cdt1-MCM2-7 bound to DNA. Nat Struct Mol Biol 20, 944–951. 10.1038/nsmb.2629.

16. Li, N., Lam, W.H., Zhai, Y., Cheng, J., Cheng, E., Zhao, Y., Gao, N., and Tye, B.K. (2018). Structure of the origin recognition complex bound to DNA replication origin. Nature 559, 217–222. 10.1038/s41586-018-0293-x.

17. Miller, T.C.R., Locke, J., Greiwe, J.F., Diffley, J.F.X., and Costa, A. (2019). Mechanism of head-to-head MCM double-hexamer formation revealed by cryo-EM. Nature 575, 704–710. 10.1038/s41586-019-1768-0.

18. Sanchez, H., Liu, Z., van Veen, E., van Laar, T., Diffley, J.F.X., and Dekker, N.H. (2023). A chromatinized origin reduces the mobility of ORC and MCM through interactions and spatial constraint. Nat Commun 14, 6735. 10.1038/s41467-023-42524-8.

19. Amasino, A.L., Gupta, S., Friedman, L.J., Gelles, J., and Bell, S.P. (2023). Regulation of replication origin licensing by ORC phosphorylation reveals a two-step mechanism for Mcm2-7 ring closing. Proc Natl Acad Sci U S A 120, e2221484120. 10.1073/pnas.2221484120.

20. Wells, J.N., Edwardes, L.V., Leber, V., Allyjaun, S., Peach, M., Tomkins, J., Kefala-Stavridi, A., Faull, S.V., Aramayo, R., Pestana, C.M., et al. (2025). Reconstitution of human DNA licensing and the structural and functional analysis of key intermediates. Nat Commun 16, 478. 10.1038/s41467-024-55772-z.

21. Remus, D., Beuron, F., Tolun, G., Griffith, J.D., Morris, E.P., and Diffley, J.F. (2009). Concerted loading of Mcm2-7 double hexamers around DNA during DNA replication origin licensing. Cell 139, 719–730. 10.1016/j.cell.2009.10.015.

22. Weissmann, F., Greiwe, J.F., Puhringer, T., Eastwood, E.L., Couves, E.C., Miller, T.C.R., Diffley, J.F.X., and Costa, A. (2024). MCM double hexamer loading visualized with human proteins. Nature 636, 499–508. 10.1038/s41586-024-08263-6.

23. Yang, R., Hunker, O., Wise, M., and Bleichert, F. (2024). Multiple mechanisms for licensing human replication origins. Nature 636, 488–498. 10.1038/s41586-024-08237-8.

24. Li, J., Dong, J., Wang, W., Yu, D., Fan, X., Hui, Y.C., Lee, C.S.K., Lam, W.H., Alary, N., Yang, Y., et al. (2023). The human pre-replication complex is an open complex. Cell 186, 98–111 e121. 10.1016/j.cell.2022.12.008.

25. Knapp, K.M., Jenkins, D.E., Sullivan, R., Harms, F.L., von Elsner, L., Ockeloen, C.W., de Munnik, S., Bongers, E., Murray, J., Pachter, N., et al. (2021). MCM complex members MCM3 and MCM7 are associated with a phenotypic spectrum from Meier-Gorlin syndrome to lipodystrophy and adrenal insufficiency. Eur J Hum Genet 29, 1110–1120. 10.1038/s41431-021-00839-4.

26. Nielsen-Dandoroff, E., Ruegg, M.S.G., and Bicknell, L.S. (2023). The expanding genetic and clinical landscape associated with Meier-Gorlin syndrome. Eur J Hum Genet 31, 859–868. 10.1038/s41431-023-01359-z.

27. Jamali, K., Kall, L., Zhang, R., Brown, A., Kimanius, D., and Scheres, S.H.W. (2024). Automated model building and protein identification in cryo-EM maps. Nature 628, 450–457. 10.1038/s41586-024-07215-4.

28. Harami, G.M., Gyimesi, M., and Kovacs, M. (2013). From keys to bulldozers: expanding roles for winged helix domains in nucleic-acid-binding proteins. Trends Biochem Sci 38, 364–371. 10.1016/j.tibs.2013.04.006.

29. Frigola, J., Remus, D., Mehanna, A., and Diffley, J.F. (2013). ATPase-dependent quality control of DNA replication origin licensing. Nature 495, 339–343. 10.1038/nature11920.

30. Nabet, B., Ferguson, F.M., Seong, B.K.A., Kuljanin, M., Leggett, A.L., Mohardt, M.L., Robichaud, A., Conway, A.S., Buckley, D.L., Mancias, J.D., et al. (2020). Rapid and direct control of target protein levels with VHL-recruiting dTAG molecules. Nature communications 11, 4687. 10.1038/s41467-020-18377-w.

31. Ge, X.Q., Jackson, D.A., and Blow, J.J. (2007). Dormant origins licensed by excess Mcm2-7 are required for human cells to survive replicative stress. Genes Dev 21, 3331–3341. 10.1101/gad.457807.

32. Ibarra, A., Schwob, E., and Mendez, J. (2008). Excess MCM proteins protect human cells from replicative stress by licensing backup origins of replication. Proc Natl Acad Sci U S A 105, 8956–8961. 10.1073/pnas.0803978105.

33. Polasek-Sedlackova, H., Miller, T.C.R., Krejci, J., Rask, M.B., and Lukas, J. (2022). Solving the MCM paradox by visualizing the scaffold of CMG helicase at active replisomes. Nat Commun 13, 6090. 10.1038/s41467-022-33887-5.

34. Alver, R.C., Chadha, G.S., and Blow, J.J. (2014). The contribution of dormant origins to genome stability: from cell biology to human genetics. DNA Repair (Amst) 19, 182–189. 10.1016/j.dnarep.2014.03.012.

35. Saxena, S., and Zou, L. (2022). Hallmarks of DNA replication stress. Mol Cell 82, 2298–2314. 10.1016/j.molcel.2022.05.004.

36. Weissmann, F., Petzold, G., VanderLinden, R., Huis In ’t Veld, P.J., Brown, N.G., Lampert, F., Westermann, S., Stark, H., Schulman, B.A., and Peters, J.M. (2016). biGBac enables rapid gene assembly for the expression of large multisubunit protein complexes. Proceedings of the National Academy of Sciences of the United States of America 113, E2564–2569. 10.1073/pnas.1604935113.

37. Zheng, G., Kanchwala, M., Xing, C., and Yu, H. (2018). MCM2-7-dependent cohesin loading during S phase promotes sister-chromatid cohesion. Elife 7. 10.7554/eLife.33920.

38. Mailand, N., and Diffley, J.F. (2005). CDKs promote DNA replication origin licensing in human cells by protecting Cdc6 from APC/C-dependent proteolysis. Cell 122, 915–926. 10.1016/j.cell.2005.08.013.

39. Zheng, S.Q., Palovcak, E., Armache, J.P., Verba, K.A., Cheng, Y., and Agard, D.A. (2017). MotionCor2: anisotropic correction of beam-induced motion for improved cryo-electron microscopy. Nature methods 14, 331–332. 10.1038/nmeth.4193.

40. Zhang, K. (2016). Gctf: Real-time CTF determination and correction. J Struct Biol 193, 1–12. 10.1016/j.jsb.2015.11.003.

41. Punjani, A., Rubinstein, J.L., Fleet, D.J., and Brubaker, M.A. (2017). cryoSPARC: algorithms for rapid unsupervised cryo-EM structure determination. Nature methods 14, 290–296. 10.1038/nmeth.4169.

42. Pettersen, E.F., Goddard, T.D., Huang, C.C., Meng, E.C., Couch, G.S., Croll, T.I., Morris, J.H., and Ferrin, T.E. (2021). UCSF ChimeraX: Structure visualization for researchers, educators, and developers. Protein Sci 30, 70–82. 10.1002/pro.3943.

43. Emsley, P., and Cowtan, K. (2004). Coot: model-building tools for molecular graphics. Acta Crystallogr. D Biol. Crystallogr. 60, 2126–2132.

44. Varadi, M., Anyango, S., Deshpande, M., Nair, S., Natassia, C., Yordanova, G., Yuan, D., Stroe, O., Wood, G., Laydon, A., et al. (2022). AlphaFold Protein Structure Database: massively expanding the structural coverage of protein-sequence space with high-accuracy models. Nucleic Acids Res 50, D439–D444. 10.1093/nar/gkab1061.

45. Afonine, P.V., Poon, B.K., Read, R.J., Sobolev, O.V., Terwilliger, T.C., Urzhumtsev, A., and Adams, P.D. (2018). Real-space refinement in PHENIX for cryo-EM and crystallography. Acta Crystallogr D Struct Biol 74, 531–544. 10.1107/S2059798318006551.

46. Davis, I.W., Leaver-Fay, A., Chen, V.B., Block, J.N., Kapral, G.J., Wang, X., Murray, L.W., Arendall, W.B., 3rd, Snoeyink, J., Richardson, J.S., and Richardson, D.C. (2007). MolProbity: all-atom contacts and structure validation for proteins and nucleic acids. Nucleic Acids Res 35, W375–383. 10.1093/nar/gkm216.

47. Mendez, J., and Stillman, B. (2000). Chromatin association of human origin recognition complex, cdc6, and minichromosome maintenance proteins during the cell cycle: assembly of prereplication complexes in late mitosis. Mol Cell Biol 20, 8602–8612. 10.1128/MCB.20.22.8602-8612.2000.

