## Supplemental Data for "Cryo-EM structure of DNA-unbound human MCM2–7 complex reveals new disease-relevant regulation"

Figures S1-S8 and Table S1

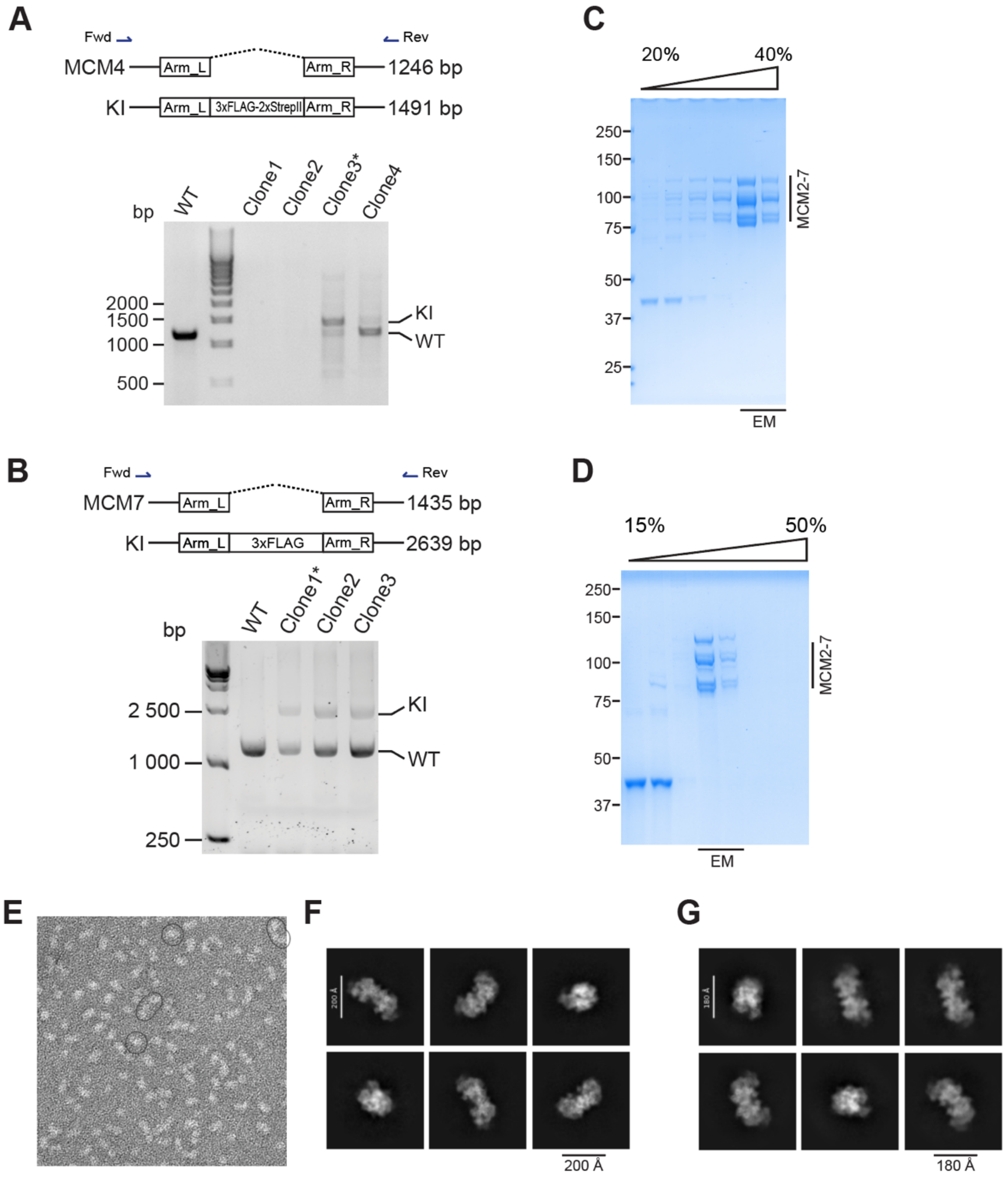


**Figure S1. Purification and negative staining EM analyses of human MCM2–7. Related to Figure 1**

(A) Genotyping of MCM4 knock-in (KI) cell lines by PCR. The clone used in this study is marked with an asterisk.

(B) Genotyping of MCM7 knock-in (KI) cell lines by PCR. The clone used in this study is marked with an asterisk.

(C) Fractionation of the MCM2–7 complex purified from MCM4 KI cells on glycerol gradient. The fractions were analyzed with Coomassie blue stained SDS-PAGE. Fractions used for cryo-EM analyses were marked.
(D) Fractionation of the MCM2–7 complex purified from MCM7 KI cells on sucrose gradient. The fractions were analyzed with Coomassie blue stained SDS-PAGE. Fractions used for cryo-EM analyses were marked.
(E) Representative negative staining TEM images of the MCM2–7 complex purified from MCM4 KI cells.
(F and G) 2D classes of MCM2–7 complexes purified from MCM4 KI (F) and MCM7 KI (G) cells.

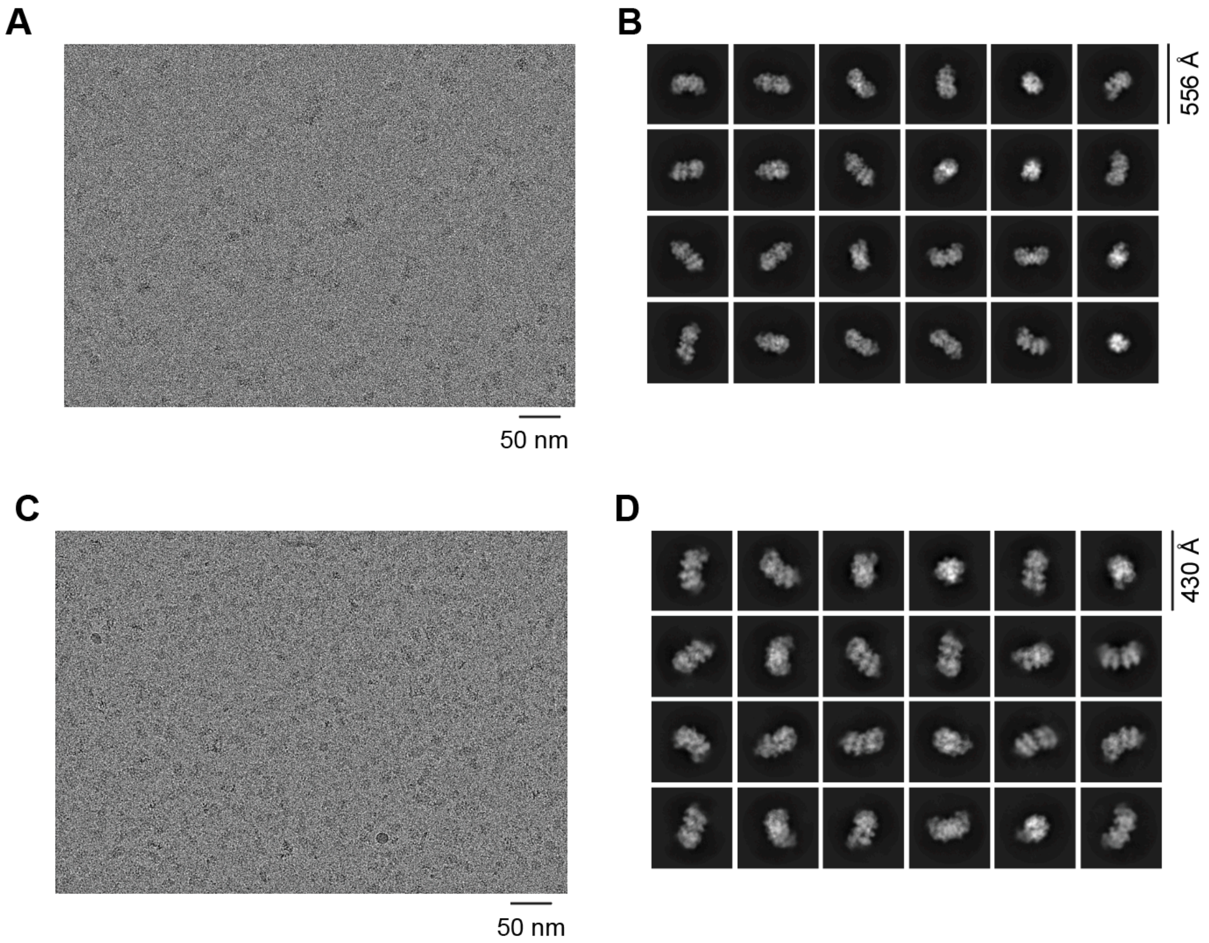


**Figure S2. Cryo-EM analysis of DNA-unbound human MCM double hexamer (DH). Related to Figure 1**

(A) Representative raw images of MCM2–7 complexes purified from MCM4 KI cells.

(B) Representative 2D class averages of MCM2–7 complexes purified from MCM4 KI cells.

(C) Representative raw images of MCM2–7 complexes purified from MCM7 KI cells.

(D) Representative 2D class averages of MCM2–7 complexes purified from MCM7 KI cells.

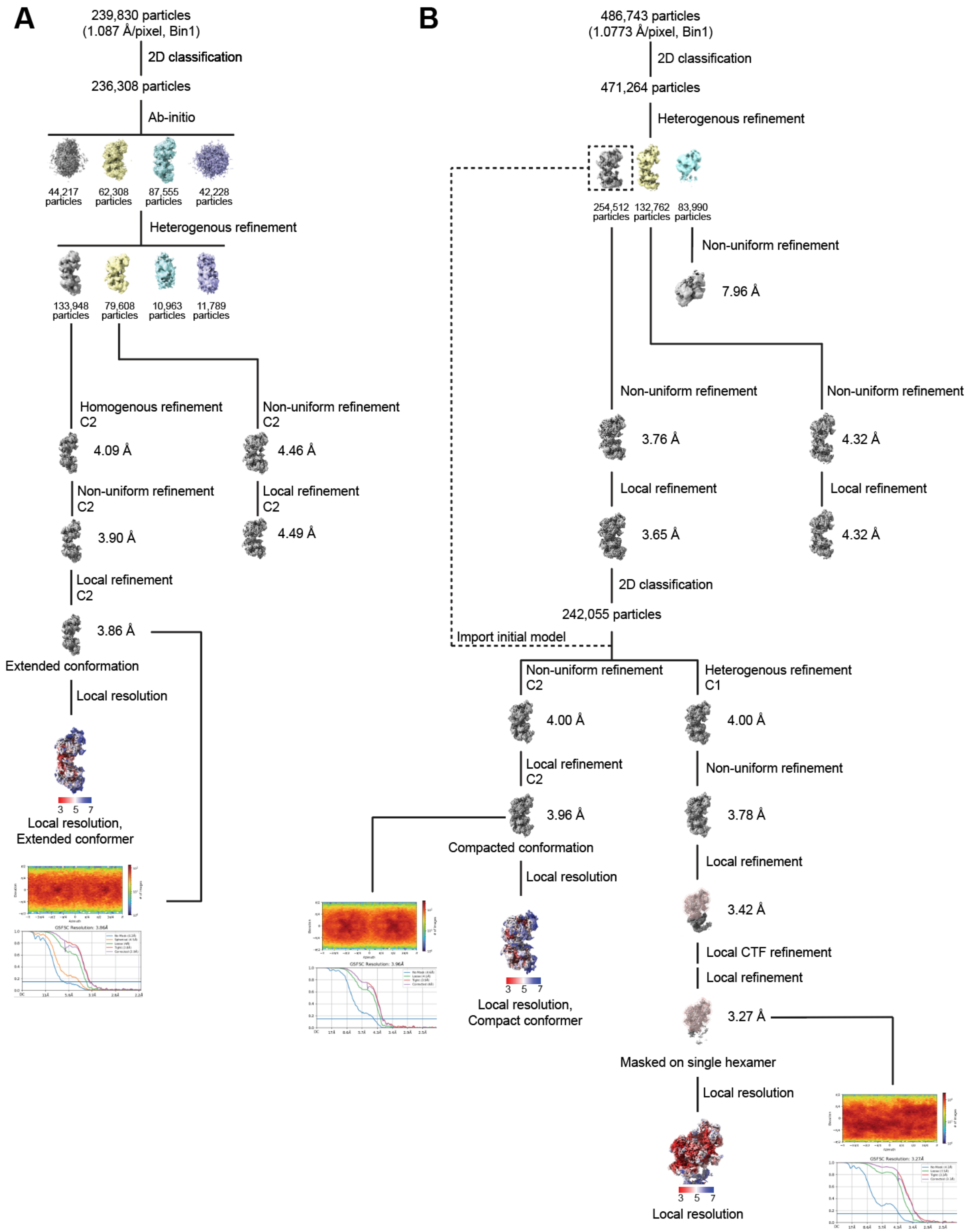


**Figure S3. Cryo-EM image processing flowchart. Related to Figure 1**

(A) Cryo-EM image processing flowchart of MCM2–7 complexes purified from MCM4 knock-in (KI) cells. Local density maps for model building and analysis in this study were generated. Particle distributions and Gold-standard Fourier shell correlation (GSFSC) of the final map were also plotted.
(B) Cryo-EM image processing flowchart of MCM2–7 complexes purified from MCM7 KI cells. Local density maps, particle distributions, and GSFSC of the final map were shown.

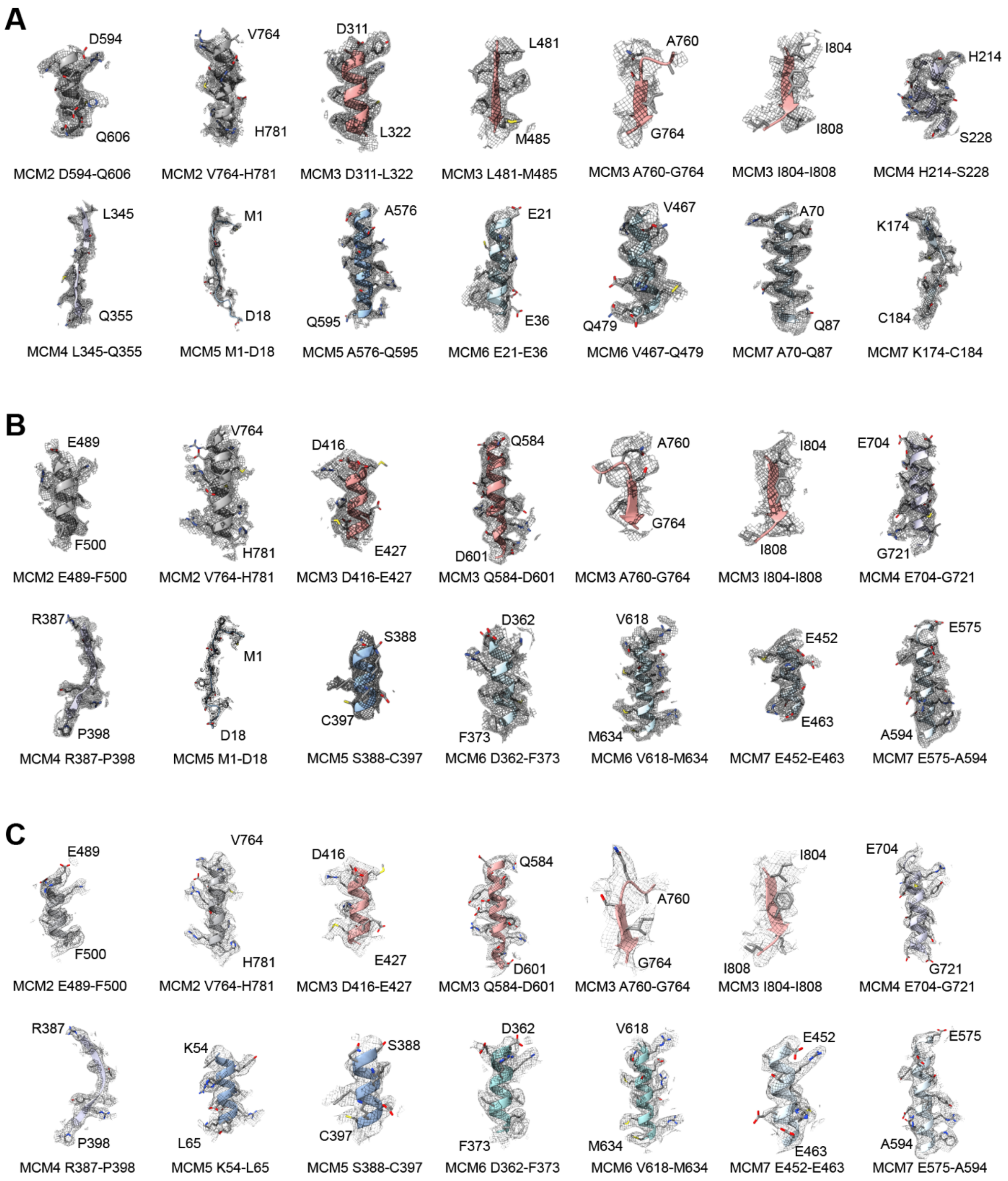


**Figure S4. Cryo-EM density maps of structural elements of DNA-unbound human MCM double hexamer (DH). Related to Figure 1**

(A) Local density maps of selected structural elements of the extended conformer of MCM2–7 DH.

(B) Local density maps of selected structural elements of the compacted conformer of MCM2–7 DH.
(C) Local density maps of selected structural elements of the MCM2–7 single hexamer (SH).

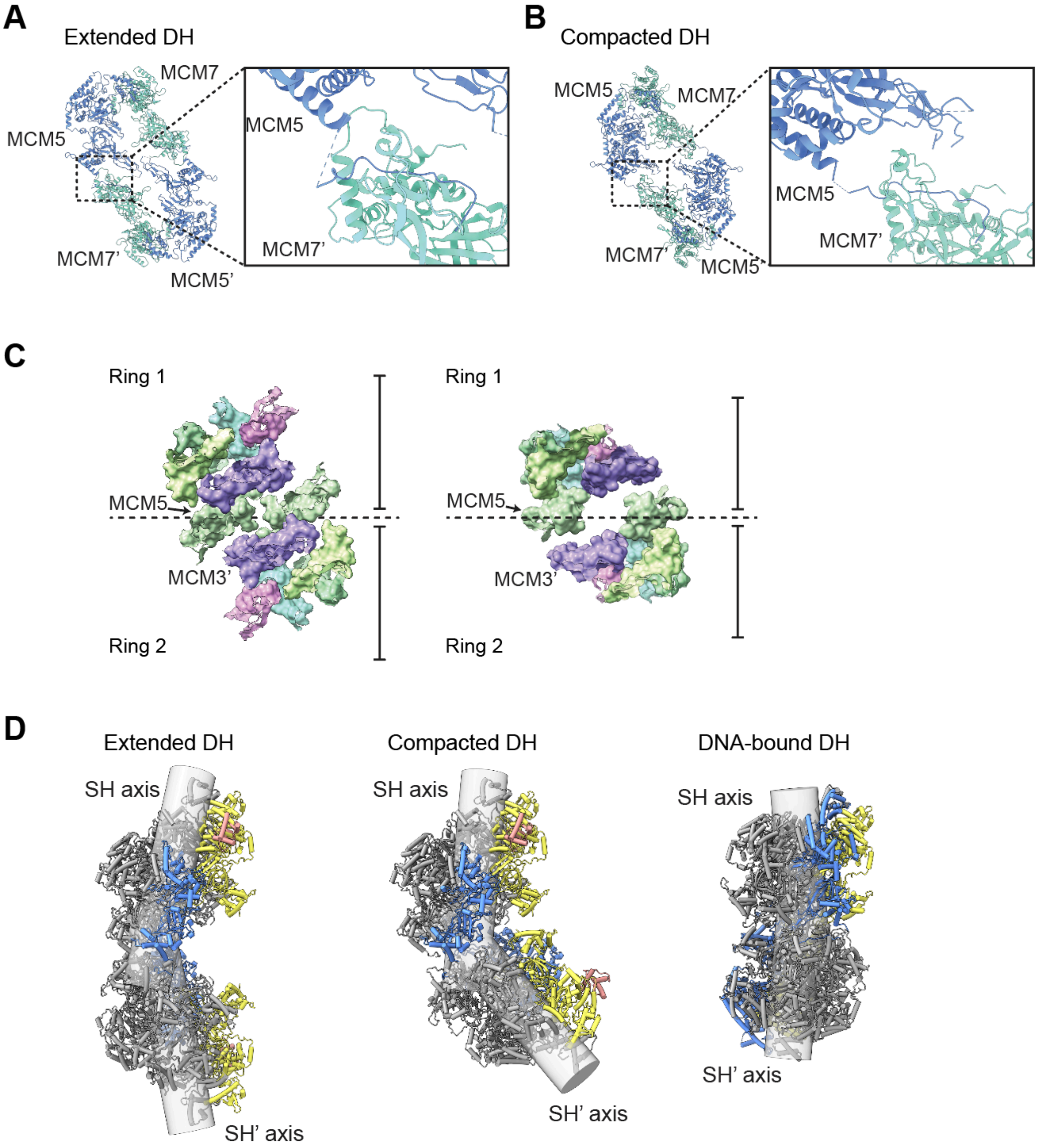


**Figure S5. The interface between two single hexamers (SH) and their central axes in the MCM2–7 double hexamer (DH). Related to Figure 1**

(A and B) Interfaces between MCM5 N-terminus extension with MCM7' in two DNA-free MCM DH conformers.

(C) back views of ZF rings in two conformations of MCM DH.

(D) Side views of open-gated extended and compact conformers of DNA-free MCM DH with MCM5 (blue), MCM2 (yellow), and MCM3 WHD (pink), in comparison with DNA-bound MCM

DH (PDB ID: 7Y1W). For DNA-free conformers, axes were defined by central channels in both SH. For DNA-bound MCM DH, axes defined by dsDNA in both SH.

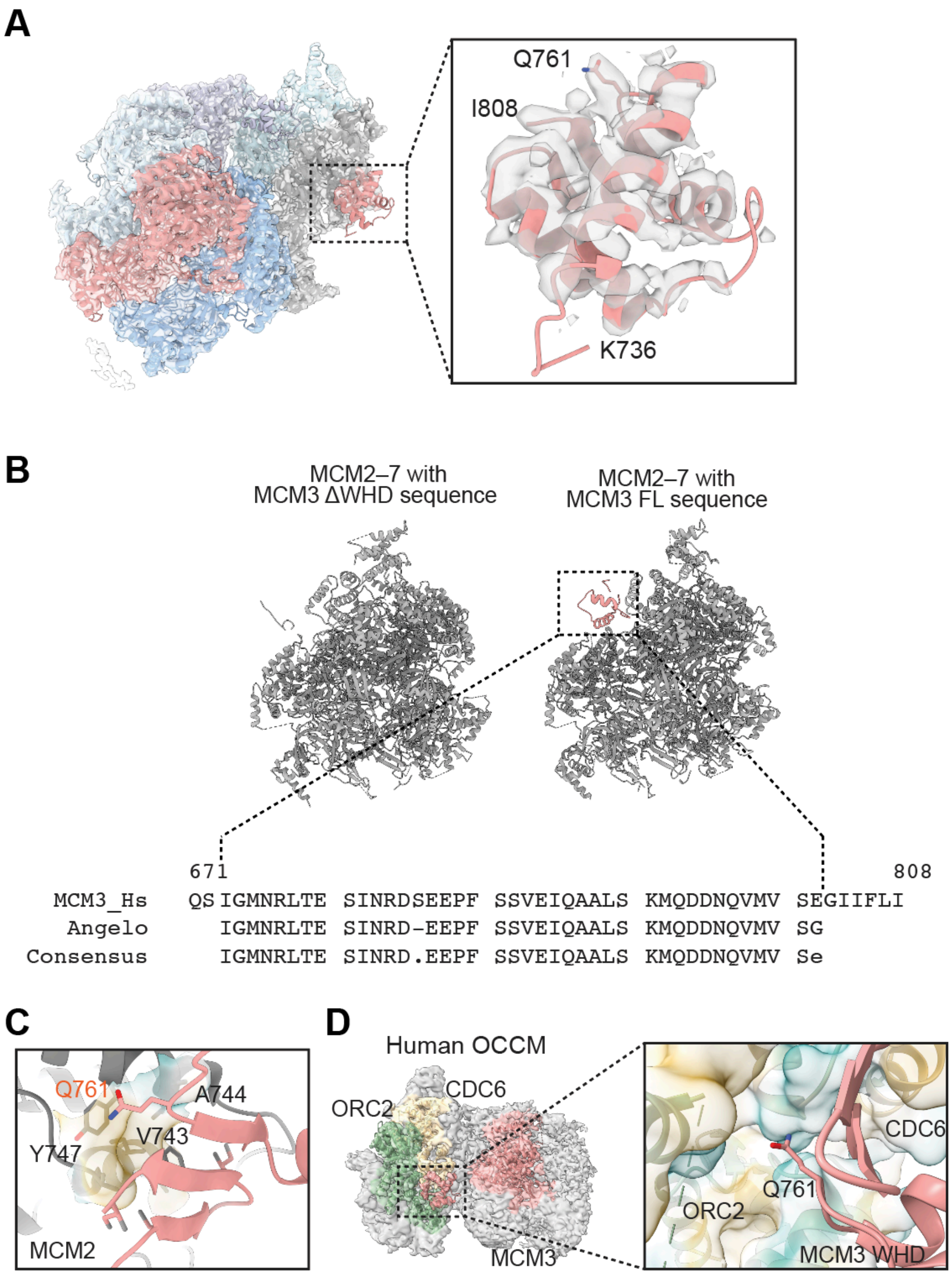


**Figure S6. MCM3 WHD binds to MCM2 in the DNA-unbound MCM2–7 DH. Related to Figure 2**

(A) Side view of the segmented cryo-EM density map of MCM2–7 SH, superimposed with the atomic model. A close-up view of MCM3 WHD is shown on the right.

(B) ModelAngelo autobuilding with inputs of the full-length amino acid sequences of MCM2–7 subunits with and without MCM3 WHD. In the full-length model, MCM3 WHD (colored pink) can be modeled in the location of the extra cryo-EM density near MCM2. An alignment between the human MCM3 WHD sequence and the sequence of autobuilt atomic model in the extra density region was shown below.

(C) The hydrophobic Q761-binding pocket on MCM2.

(D) Cryo-EM density map of human OCCM superimposed with the atomic model of ORC2 (green), CDC6 (yellow), and MCM3 (pink). A close-up view of the ORC2-CDC6-MCM3 WHD interface is shown on the right.

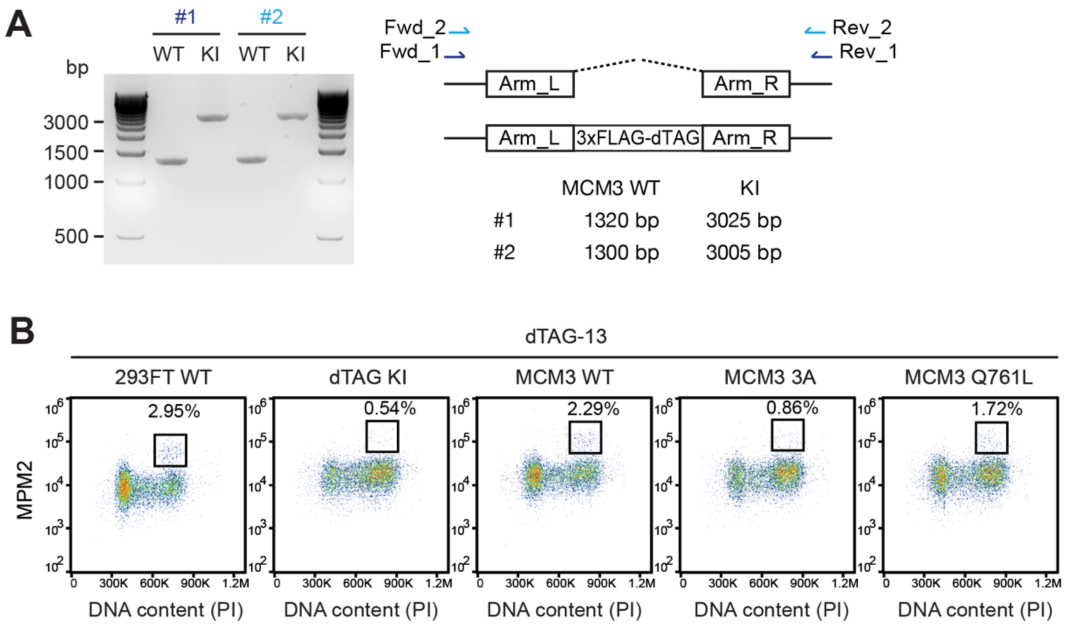


**Figure S7. MCM3 WHD mutations induce ATR-CHK1-dependent G2 arrest in human cells. Related to Figure 3**

(A) Genotyping of MCM3 knock-in (KI) cells by PCR.

(B) Representative flow cytometry plots of the indicated 293FT cells treated with dTAG-13 (1 µM) for 16 h. 293FT wild type (WT), the parental cell line without gene editing; dTAG KI, 293FT cells with dTAG KI at the endogenous *MCM3* loci; MCM3 WT, dTAG KI 293FT cells stably expressing HA-MCM3 wild type; MCM3 3A, dTAG KI 293FT cells stably expressing HA-MCM3 3A; MCM3 Q761L, dTAG KI 293FT cells stably expressing HA-MCM3 Q761L. The mitotic cells (MPM2-positive cells with 4N DNA content) are boxed. Mitotic indices are indicated.

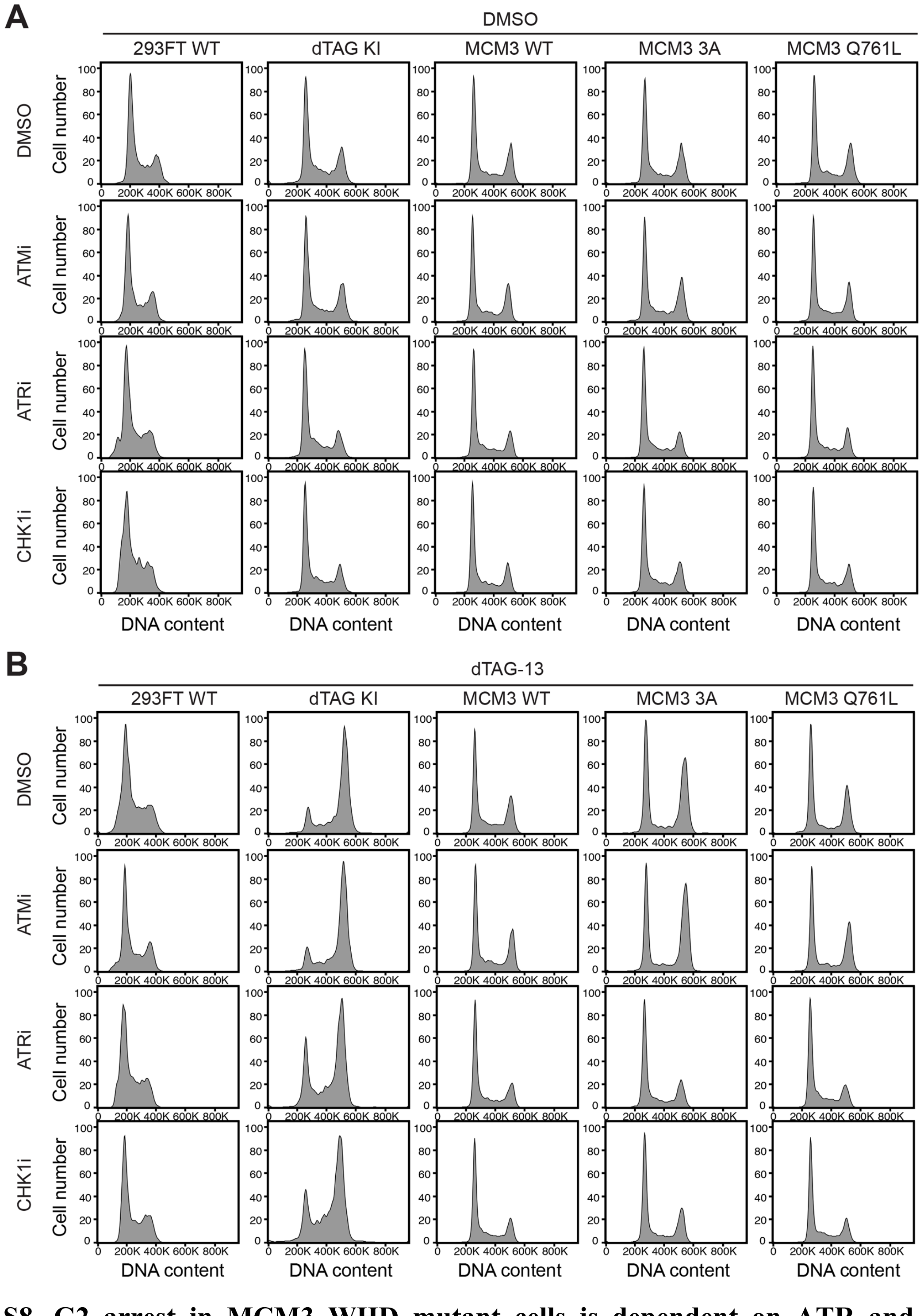


**Figure S8. G2 arrest in MCM3 WHD mutant cells is dependent on ATR and CHK1. Related to Figure 3**

(A) Representative FACS histograms of indicated 293FT cells treated with DMSO and the indicated chemical inhibitors. 293FT wild type (WT), the parental cell line without gene editing; dTAG KI, 293FT cells with dTAG KI at the endogenous *MCM3* loci; MCM3 WT, dTAG KI

293FT cells stably expressing HA-MCM3 wild type; MCM3 3A, dTAG KI 293FT cells stably expressing HA-MCM3 3A; MCM3 Q761L, dTAG KI 293FT cells stably expressing HA-MCM3 Q761L.

(B) Representative FACS histograms of indicated 293FT cells treated with dTAG-13 and the indicated chemical inhibitors.

### Table S1. Cryo-EM data collection, refinement, and validation statistics

| | Endogenous MCM2–7 | | | Recombinant MCM2–7 | |
|---|---|---|---|---|---|
| | Extended DH | Compacted DH | SH, masked | WT | 3A |
| **Data Collection and Processing** | | | | | |
| Microscope | FEI Titan Krios | | | | |
| Magnification | 81,000 | | | | |
| Voltage | 300 | | | | |
| Electron dose (e$^-$/Å$^2$) | 50 | | | | |
| Detector | Gatan K3 Summit | | | | |
| Defocus range (um) | -1.0 to -2.0 | | | | |
| Pixel size (Å/pixel) | 1.087 | 1.0773 | | | |
| Micrographs (no.) | 5,178 | 3,662 | | 5,101 | 4,010 |
| Initial particles (no.) | 239,830 | 486,743 | | 1,658,759 | 5,985,753 |
| Final particles (no.) | 133,948 | 242,055 | | 184,028 | 111323 |
| Symmetry imposed | C2 | | C1 | | |
| Map resolution (Å) | 3.86 | 3.96 | 3.27 | 3.19 | 3.80 |
| FSC threshold | 0.143 | | | | |
| **Model composition** | | | | | |
| Protein residues | 8316 | 8316 | 4139 | / | / |
| Ligands | ATP: 4<br>ADP: 2 | ATP: 2<br>ADP: 8 | ATP: 1<br>ADP: 4 | | |
| **Refinement** | | | | | |
| Initial model used | AlphaFold3 | | | | |
| Model resolution (Å) | 3.8 | 3.9 | 3.2 | | |
| FSC threshold | 0.143 | 0.143 | 0.143 | | |
| Map sharpening B factor (Å$^2$) | 112.8 | 140.9 | 113.8 | | |
| **Validation** | | | | | |
| MolProbility score | 2.34 | 2.36 | 2.26 | | |
| Clash score | 12.88 | 12.71 | 11.3 | | |
| Rotamer outliers (%) | 1.98 | 1.98 | 1.63 | | |
| $C_b$ outliers (%) | 0 | 0 | 0.03 | | |
| Bonds length (Å) | 0.003 | 0.003 | 0.003 | | |
| Bonds Angle (°) | 0.67 | 0.649 | 0.649 | | |
| **Ramachandran plot (%)** | | | | | |
| Favored | 91.9 | 90.9 | 90.62 | | |
| Allowed | 7.95 | 8.95 | 9.04 | | |
| Outliers | 0.15 | 0.15 | 0.34 | | |
